# Longitudinal impacts of habitat fragmentation on *Bartonella* and hemotropic *Mycoplasma* dynamics in vampire bats

**DOI:** 10.1101/2025.07.21.665613

**Authors:** Lauren R. Lock, Kristin E. Dyer, Dmitriy V. Volokhov, Anni Yang, M. Brock Fenton, Nancy B. Simmons, Daniel J. Becker

## Abstract

Habitat fragmentation can have negative impacts on wildlife including increased risk of infectious disease. To assess spatiotemporal changes in pathogen dynamics in vampire bats (*Desmodus rotundus*) in response to habitat fragmentation, we used general linear mixed models to investigate the influence of site, year, and tree cover on the prevalence of *Bartonella* and hemotropic *Mycoplasma* (hemoplasma) in bats in one large and one small forest fragment in northern Belize across seven years. *Bartonella* was marginally more prevalent in later years, while year and site differences in hemoplasma infections were driven by a peak in prevalence in the third year of the study in the small fragment. *Bartonella* prevalence increased with forest loss, but only in the large fragment, whereas hemoplasma prevalence showed a marginal negative response to forest loss. The effects of site, year, and forest loss on infection likelihood varied by pathogen genotype. Neither site nor year affected *Bartonella* genotypes, but one genotype was positively associated with tree cover. Two hemoplasma genotypes were influenced by year, but with differing trends. One genotype increased with tree cover regardless of site while another increased with forest loss at the small fragment only. Our work demonstrates that the effects of habitat fragmentation on infection prevalence depended on both the pathogen and specific genotype. Our findings complicate expectations of how habitat fragmentation affects infectious disease dynamics in bats. As such, management practices aimed at mitigating the impacts of infectious diseases in fragmented systems should be tailored to specific pathogens of concern.

## Introduction

Habitat fragmentation is a process in which an undisturbed habitat is broken into multiple patches. Because it is almost always coupled with habitat loss, it is difficult to differentiate between effects that are due to loss and effects due to fragmentation (Laurance, 2008; Didham et al., 2012). While no two ecosystems or species are exactly alike in their response to fragmentation, most often impacts on biodiversity are detrimental (Haddad et al., 2015). Island Biogeography Theory (IBT) is used commonly to predict the effects of habitat fragmentation on biodiversity (Laurance, 2008). According to IBT, when considering a habitat patch to be analogous to an island, diversity is expected to be lower in small, isolated patches compared to large, easily accessible patches (MacArthur & Wilson, 1967). Island Biogeography Theory can also be extended to predict the effects that habitat fragmentation has on infectious disease dynamics (Reperant, 2010; Shaw et al., 2024). This application deviates from the typical expectations of IBT – pathogens should be able to capitalize on the effects that fragmentation has on hosts, allowing pathogens to more easily invade small, more isolated habitat patches.

However, true outcomes are complicated by additional considerations centered around hosts and their responses to fragmentation (i.e. alterations to host richness, abundance, and density; host immunity; host matrix use; Faust et al., 2018; Becker et al., 2020a).

The tropical forests of the world have been subjected to exceptionally high rates of habitat fragmentation compared to other ecoregions since 2000 (Ma et al., 2023). In the Neotropics, which are considered to be the most biodiverse region on Earth, this degradation imperils an area containing at least one-third of global biodiversity (Raven et al., 2020) and eight of the world’s 34 biodiversity hotspots (Mittermeier et al., 2011). Additionally, continued fragmentation risks the increase of infectious disease emergence from wildlife (Plowright et al., 2021; Gibb et al., 2024).

Bats (order: Chiroptera) are popular and epidemiologically relevant models for studying infectious disease dynamics in wildlife. With over 1,480 species spread across all continents except Antarctica, bats are diverse, widely distributed, and make up the second largest mammalian order next to rodents (order: Rodentia) (Fenton & Simmons, 2020; Simmons & Cirranello, 2025). Bats are known reservoirs of a number of zoonotic pathogens, including but not limited to rabies, Nipah, and Hendra viruses (Banyard et al., 2011; Halpin et al., 2011; Letko et al., 2020). In addition to their natural habitats, many bat species are able to live in close proximity to humans, roosting in houses, bridges, and other anthropogenic structures, offering opportunities for pathogen transmission between bats and humans and/or domestic livestock and pets (Streicker et al., 2013; Jackson et al., 2023; Betke et al., 2024). As the only mammal capable of flight, bats also have the potential to disperse pathogens across moderate to long distances (Moussy et al., 2013; Plowright et al., 2015; Voight et al., 2017).

The common vampire bat (*Desmodus rotundus*), which is prevalent throughout the Neotropics and often occurs in abundance in areas that have been disturbed for agricultural purposes, is of particular interest to studies of zoonotic pathogens and habitat fragmentation. As vampire bats are dietary specialists that subsist entirely on blood (primarily on that of mammals), the species thrives in close proximity to livestock, especially cattle and pigs (Turner 1975; Delpietro et al., 1992; Bobrowiec et al., 2015). This leads to an increased potential for contact with domestic animals and humans compared to other bat species. Vampire bats are highly social and aggregate in colonies of that range in size from a dozen to hundreds of individuals (Trajano et al., 1996; Streicker et al., 2012), creating ample opportunity for density-dependent pathogen transmission between bats (Lloyd-Smith et al., 2005). Their social interactions include behaviors that can facilitate contact-driven pathogen transmission, such as allogrooming, aggressive encounters between males, a high degree of parental care in females, and “friendships” between bats that often involve the sharing of regurgitated blood meals (Wilkinson 1984, Carter, 2024). Vampire bat home ranges vary from ∼2 km to 5 km (Burns & Crespo, 1975; Trajano, 1996) but movements of up to 54 km have also been recorded (Delpietro et al., 2017), indicating the potential for pathogen dispersal across moderate distances.

While much of the research pertaining to infectious diseases in vampire bats focuses on rabies virus due to the potential for transmission to humans or domestic animals via blood feeding (Heckley et al., 2023), vampire bats also are known to carry other blood-borne pathogens such as *Bartonella* spp. (Becker at al., 2018) and hemotropic mycoplasmas (hemoplasmas) (Volokhov et al., 2017). Both are bacterial pathogens thought to be transmitted through a combination of direct and vector-borne contact. Although efforts to establish the primary mode of transmission for each pathogen are ongoing, current evidence suggests that *Bartonella* is spread primarily through arthropod vectors (Judson et al., 2015; Becker et al., 2018a) while bat-to-bat contact is likely the predominant driver of hemoplasma transmission (Willi et al., 2007).

While both infections are likely facilitated by the species’ highly social nature and by parasitism by bat flies and fleas (Judson et al., 2015; Volokhov et al., 2017), differences in their transmission likely mean that bartonellae and hemoplasmas could also differ in their responses to habitat fragmentation.

Bartonellae and hemoplasmas are most often studied in vampire bats in the context of zoonoses, as some lineages can infect humans. For example, *Candidatus* Bartonella mayotimonensis has been detected in several bat species (Veikkolainen et al., 2014; Lilley et al., 2017) and implicated in human cases of endocarditis (Lin et al., 2010), while *Candidatus* Mycoplasma haematohominis, reported to cause hemolytic anemia in humans (Steer et al., 2011), shares high genetic similarity to the 16S rRNA gene from hemoplasmas detected in Neotropical bats sympatric with vampire bats (Volokhov et al., 2023). However, many *Bartonella* and hemoplasma genotypes are considered to be host specific (Pitcher & Nicholas, 2005; Frank et al., 2018) and are sufficiently prevalent within vampire bat populations to make them ideal for studying the effects of habitat fragmentation on infectious disease dynamics in wildlife (Volokhov et al., 2017; Becker et al., 2018a).

Because habitat fragmentation is an ongoing process and its effects are often not immediately observable, longitudinal studies are needed to capture patterns in infectious disease dynamics over time (Heckley et al., 2023). To assess temporal changes in pathogen dynamics in bats impacted by habitat fragmentation, we compared the prevalence of all bartonellae and hemoplasmas as well as that of each pathogen genotype, between vampire bats in two habitat patches within a fragmented forest system in northern Belize across seven years. We also quantified the amount of tree cover lost within the area between and surrounding both habitats (hereafter referred to as the matrix) during the last five years of this study (land use/land cover data were not available for the first two years). Using this information, we tested the following hypotheses: infections will be more prevalent in 1) the smaller habitat patch and in later years in response to increased anthropogenic habitat fragmentation; 2) reproductive females than in males or non-reproductive females due to trade-offs between support for immune functions and support for lactation and pregnancy; and 3) subadults than in adult bats that may have acquired immunity. Additionally, we expected that the direction or magnitude of effect would vary by genotype within each pathogen.

## Methods

### Study Areas

Sample collection took place in the Orange Walk District of Belize, specifically within a preserved forest, the Lamanai Archeological Reserve (LAR; 17.76512, 88.65200), and a preserved forest fragment, Ka’Kabish (KK; 17.81455, 88.73075) (Fig. 1). The LAR is a ∼450 ha semi-deciduous forest composed mostly of secondary growth within a protected reserve.

**Figure 1.**
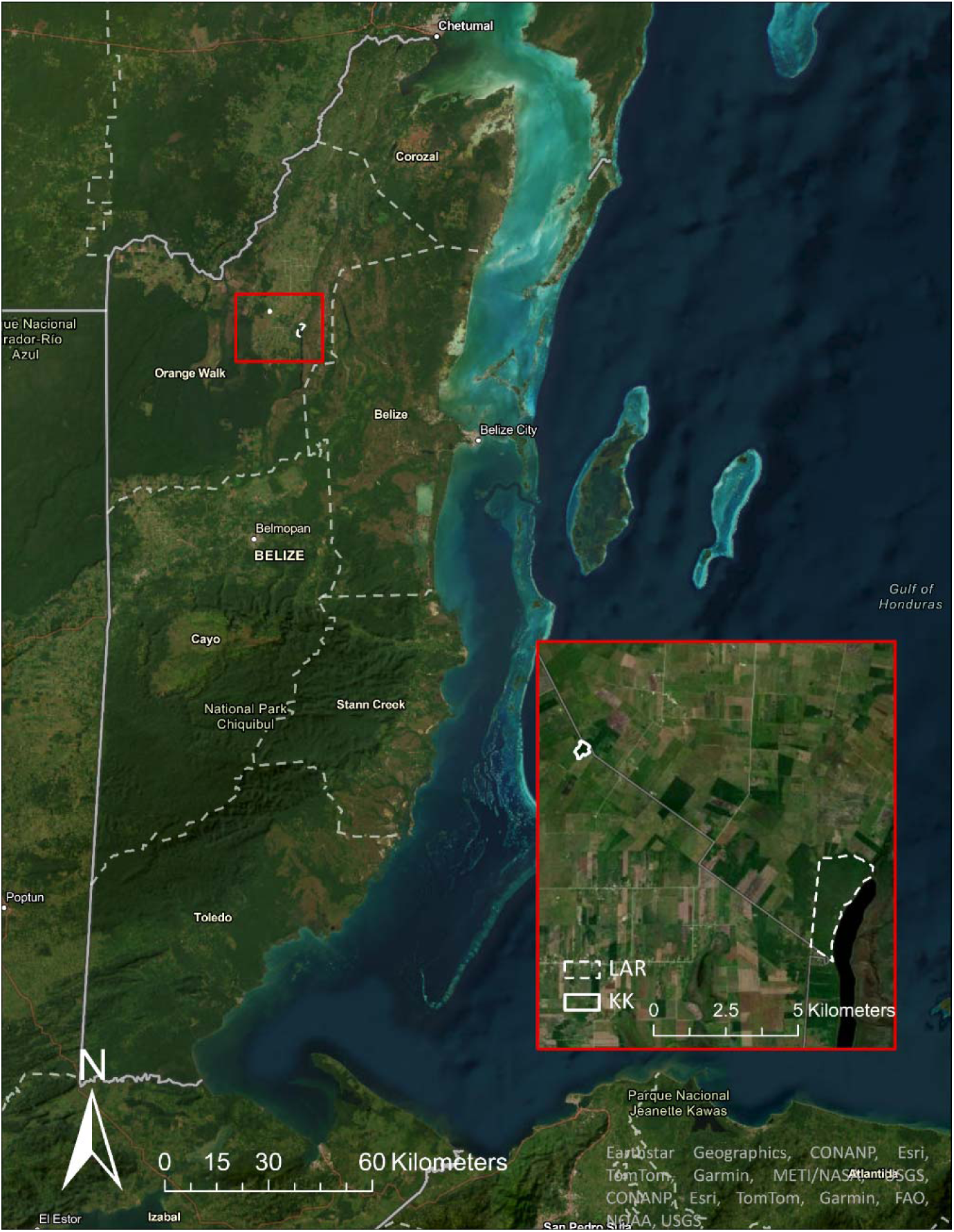
Location and land cover of the Lamanai Archaeological Reserve (LAR) and Ka’Kabish (KK) bat sampling sites in the Orange Walk District of Belize.

Ka’Kabish is a ∼45 ha remnant forest fragment bordered by agricultural fields. These sites were connected historically by contiguous forest, but they have since been separated by roughly 10 km of forest fragments and cleared land over about the last twenty years (Becker et al., 2021; Herrera et al., 2018; Ingala et al., 2019).

### Bat sampling

Vampire bats were captured using mist nets over a period of 6–13 nights during each year (6 nights/year from 2015–2017, 13 nights/year during 2018, 2019, 2021, and 2022). Samples were collected in April and May of each year except for 2020 and 2021. Sampling did not occur in 2020 due to the COVID-19 pandemic but was resumed in November 2021. Samples were not collected from KK in 2021 due to the abandonment of a major roost by the resident bats.

Vampire bats were captured at this site in the subsequent year but at lower numbers. After capture all bats were held in individual clean cloth bags until processing. We recorded the sex, reproductive status, age class, weight, forearm length, and the unique identification number of each bat prior to sampling. We took blood samples by lancing the propatagial vein using 23-gauge sterile needles and collecting blood in heparinized capillary tubes, which was then transferred to Whatman FTA cards. Samples were stored and shipped at ambient temperature until long-term storage at-20 °C. We marked each bat with an incoloy wing band with a unique alphanumeric code (Porzana Inc.) prior to release.

All fieldwork was conducted following guidelines for the safe and humane handling of bats published by of the American Society of Mammalogists (Sikes 2016) and was approved by the Institutional Animal Care and Use Committees of the University of Georgia (A2014 04-016-Y3-A5), American Museum of Natural History (AMNHIACUC-20170403, AMNHIACUC-20180123, AMNHIACUC-20190129), and University of Oklahoma (2022–0197). Fieldwork and sampling were authorized by the Belize Forest Department under permits CD/60/3/15(21), WL/1/1/16(17), WL/2/1/17(16), WL/2/1/17(19), WL/2/1/18(16), FD/WL/1/19(06), FD/WL/1/19(09), and FD/WL/1/21(12).

### Tree cover quantification

Because of the difficulties associated with differentiating between the effects of habitat loss and habitat fragmentation (Laurance, 2008; Didham et al., 2012), we used the change in tree cover in the matrix surrounding the LAR and KK over time as a proxy for habitat fragmentation. We quantified land-use change between sampling years with ArcGIS Pro version 3.0.3 (Esri Inc.) using data from the Sentinel-2 10m Land Use/Land Cover Time Series produced by Impact Observatory, Microsoft, and Esri (Karra, Kontgis, et al. 2021). Using the midpoint between the LAR and KK as the centroid, we cropped the map to a 10 km extent to encompass an estimated 5 km home range of bats for each site based on previous estimates of Neotropical bat movement patterns (Trajano, 1996; Fenton et al., 2000; Fleming et al., 1972; Becker et al., 2021). We masked the extents of the LAR and KK in order to focus our analysis on the matrix. Individual binary layers were then created for each available year (2017–2022) in which pixels belonging to the “Trees” land cover classification were assigned a value of 1, while pixels belonging to all other land cover classifications were assigned a value of 0. We quantified the change in tree cover from one year to the next for sequential pairs of years by subtracting the binary layer extent from each pair of successive years (i.e. 2018 - 2017, 2019 - 2018, etc.). The number of cells with a value of 1 and the number of cells with a value of-1 in the resulting layer was each multiplied by the pixel size of the map (10 m^2^) and converted to square kilometers to determine the area gained and lost, respectively. The area lost was subtracted from the area gained to determine the net change in area per year. We calculated the percent change in forest cover per year by dividing the net change in area by the total area in the first year of each pair of years.

### Pathogen diagnostics and genotyping

Genomic DNA was extracted from 431 blood samples (319 from LAR and 112 from KK) using Qiagen QIAamp DNA Investigator Kits. Extracted DNA was screened for *Bartonella* spp. through nested PCR amplification of the partial citrate synthase (*gltA*) gene (Norman et al., 1995; Birtles & Raoult, 1996). All PCRs used blank FTA card punches and ultrapure water as extraction and negative controls, respectively. Results were visualized using electrophoresis of a 2% agarose gel. We did not include positive controls of *Bartonella* spp., due to cross-contamination risks from nested PCR, and amplicons of expected size (∼300 bp) were identified during gel electrophoresis. These positive amplicons were cleaned using Zymo DNA Clean and Concentrator-5, Zymoclean Gel DNA Recovery Kits, or Applied Biosystems ExoSAP-IT™ PCR Product Cleanup Reagent. Cleaned amplicons were Sanger sequenced at the Georgia Genomics Facility (2015 and 2016 samples) and the North Carolina State University Genomics Sciences Laboratory (2017–2019, 2021, and 2022 samples). DNA extracts were also screened for *Mycoplasma* spp. through PCR amplification of the partial 16S ribosomal RNA (rRNA) genes (Volokhov et al., 2011). For a subset of 16S rRNA–positive samples, we also attempted to amplify the partial 23S ribosomal RNA (rRNA) gene and/or the RNA polymerase beta subunit (*rpoB*) gene to build a broader library of these genes for hemoplasmas (Table S1; Volokhov et al., 2011; Becker et al., 2025). *Candidatus* Mycoplasma haematozalophi DNA was used as a positive control. Hemoplasma amplicons were Sanger sequenced at Psomagen.

Quality control of *Bartonella* sequences was performed by manually trimming and editing in Geneious version 2023.2.1. Hemoplasma 16S rRNA sequences were screened for chimeric sequences using DECIPHER (Wright et al., 2012) and UCHIME (Edgar et al., 2011). Related sequences were identified using NCBI BLASTn, and the top hits were aligned with our sequences and reference sequences using MUSCLE, also in Geneious. Phylogenetic analysis was performed in NGPhylogeny.fr (Lemoine et al., 2019) using maximum likelihood with smart model selection (PhyML + SMS). Sequences were assigned to genotypes based on phylogenetic similarity using previously established cutoffs for the *gltA* gene for bartonellae (96%; La Scola et al., 2003; Becker et al., 2018a) and the 16S rRNA gene for hemoplasmas (98.5%; Becker et al. 2020b). Data from 2015 and 2016 were published previously (Volokhov et al., 2017; Becker et al., 2018a), as was a subset of vampire bat data focused only on the LAR from 2017, 2018, and 2019 (DeAnglis et al., 2024). We deposited 199 new sequences to GenBank (*Bartonella* accession numbers PQ758803–PQ759000, PV067250; see Table S1 for hemoplasma accession numbers). Because we amplified multiple hemoplasma genes for a subset of individual bats, we also use these multi-loci data here to propose novel *Candidatus* species when the same genotype was identified in at least two samples using 16S rRNA and at least one other marker (i.e., 16S rRNA and 23S rRNA; or 16S rRNA, 23S rRNA, and *rpoB*) (Volokhov et al., 2012, 2023).

### Statistical analysis

We used the *prevalence* package (Devleesschauwer et al., 2022) in R version 4.2.1 (R Core Team, 2022) to estimate site-and year-specific infection prevalence per pathogen including corresponding 95% confidence intervals. We then used a cross-correlation analysis of the prevalence time series per site to quantify the synchrony in infection dynamics (Shumway & Stoffer, 2000). Next, we used generalized linear mixed models (GLMMs) with the *lme4* package (Bates et al., 2015) to test our hypotheses about habitat fragmentation and temporal changes in infection dynamics. We included bat identification number (hereafter bat ID) as a random effect in order to account for recaptured bats that were sampled multiple times across the study period (n = 59). We fit an initial suite of four GLMMs separately for bartonellae and for hemoplasmas to first test if the effect of capture site (the LAR and KK) varied with year to influence the likelihood of infection while also identifying additional predictors such as age class, sex, and reproductive status. We then used four similar GLMMs to test if regional forest loss per year or site-specific factors influenced the likelihood of infection; year-specific area of tree cover in the matrix was used as a fixed effect in place of the year variable. As land cover data were only available from 2017 onwards, data from bats captured in 2015 and 2016 were excluded from this analysis (*n* = 358). All GLMMs used a binomial distribution and the nlminb optimizer. For each pathogen, we used corrected Akaike information criterion (AICc) to compare each four-model suite that considered additive and interactive effects of site, year or tree cover, and individual-level covariates using the *MuMIn* package (Burnham & Anderson, 2002). We computed Akaike weights (*w_i_*) within each model suite and considered models within two ΔAIC of the top model to be competitive. We also calculated marginal and conditional *R^2^* (*R^2^* and *R^2^*) using the *performance* package (Lüdecke et al., 2021).

To assess pathogen diversity over time, we used a Chi-squared test to compare the frequencies of each *Bartonella* and hemoplasma genotype among and within years. We calculated frequencies for both sites pooled together for each year since there were no samples for KK in 2021 and only three positive samples for KK in 2022. For any genotype with at least 10 observations, we also tested effects of site, year, and tree cover using the same modeling approach used for infection status above. We created new binary infection status variables for each genotype for bats only infected with bartonellae or hemoplasmas, respectively. We then used AICc to compare models within each suite of GLMMs. For simplicity, bats identified as infected with more than one genotype were classified according to the genotype most clearly distinguishable from the sequence chromatogram.

Lastly, we used separate general linear models (GLMs) with a Poisson distribution per pathogen to determine if the total number of genotypes observed infecting a recaptured individual bat across the study duration was influenced by sex, number of captures, minimum age, and whether a bat switched from subadult to adult during its capture history. We also used binomial GLMs to determine if the same fixed effects influenced the likelihood a recaptured bat would switch infection status or genotype.

## Results

Across all years, 233 bats tested positive for *Bartonella* infections out of 427 total bats tested, and 241 bats tested positive for hemoplasma infections out of 430 total bats tested. Throughout the seven-year study period, *Bartonella* prevalence ranged from 36.4% (CI = 0.15–0.65) to 77.7% (CI = 0.59–0.89) in the LAR (∼450 ha) and 30.0% (CI = 0.11–0.60) to 66.7% (CI = 0.35–0.88) in KK (∼45 ha). At the same time, hemoplasma prevalence ranged from 47.8% (CI = 0.34– 0.62) to 81.8% (CI = 0.52–0.95) in the LAR and 33.3% (CI = 0.12–0.65) to 90.5% (CI = 0.71–0.97) in KK (45 ha) (Fig. 2A-B). Cross-correlation analysis suggested high synchrony in prevalence across annual lags in *Bartonella* infections, while we only observed high synchrony in prevalence at the negative two-year lag for hemoplasma infections, with weak synchrony in prevalence at all other annual lags (Fig. 2C-D).

**Figure 2.**
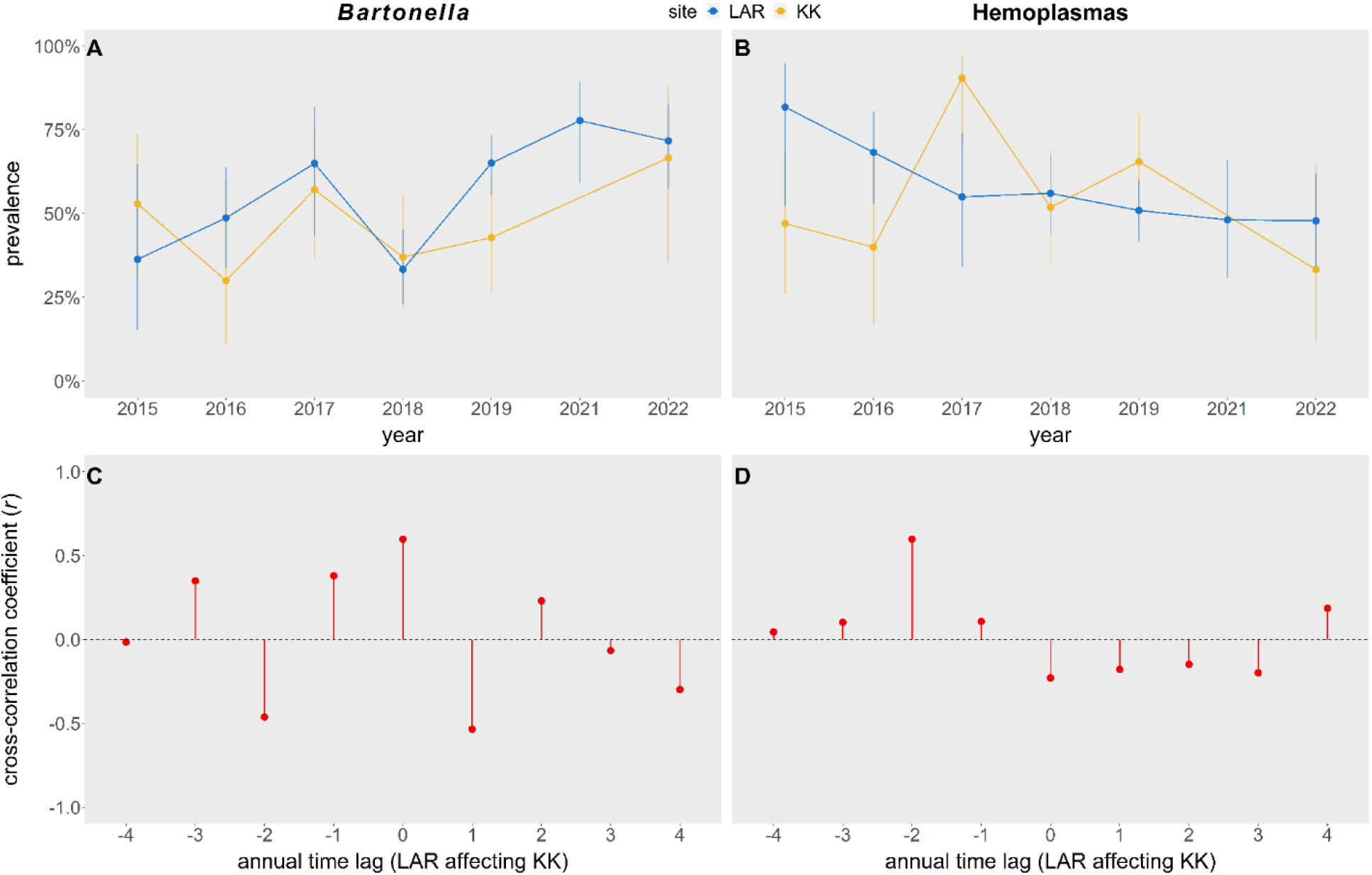
*Bartonella* (A) and hemoplasma (B) prevalence by year in the LAR (blue) and KK (yellow). Point estimates are shown with 95% confidence intervals (Wilson interval). Cross-correlation analysis of *Bartonella* (C) and hemoplasma (D) prevalence in the LAR affecting that in KK.

The forest cover in the 10 km of matrix around the central point between the LAR and KK was reduced by 23.8% from 2017-2022. Most forest loss occurred within the first two years of the timespan, with an 8.2% decrease from 9.7 km^2^ in 2017 to 8.9 km^2^ in 2018, then another 14.5% decrease to 7.6 km^2^ in 2019. Forest cover then remained relatively stable through the end of the study, rising briefly by 3.6% to 7.8 km^2^ in 2020 before falling again by 4% to 7.5 km^2^ in 2021 and by another 2.3% to 7.4 km^2^ in 2022 (Fig. S1).

### Spatiotemporal predictors of Bartonella infection

Among four GLMMs predicting *Bartonella* infection status across all years (2015–2022), the most competitive model included additive effects of site, year, sex, reproductive status, and age class (Table 1). Infection was best predicted by year (Table S2), with bats having a significantly different likelihood of infection across pairwise year comparisons. Bats had a marginally higher likelihood of being infected in 2021 (OR = 0.22, p = 0.06) and 2022 (OR = 0.30, p = 0.06) compared to 2015. Bats were also marginally more likely to be infected in 2019 (OR = 0.41, p = 0.06) and significantly more likely to be infected in 2021 (OR = 0.19, p = 0.02) and 2022 (OR = 0.26; p = 0.02) compared to 2016. The likelihood of infection was marginally higher in 2017 (OR = 2.82, p = 0.06) compared to 2018 and significantly lower in 2018 compared to 2019 (OR = 0.35, p < 0.01), 2021 (OR = 0.16, p < 0.01), and 2022 (OR = 0.22, p < 0.01) (Fig. 2A). Sex and reproductive status also predicted *Bartonella* infection, with males having a significantly higher likelihood of infection than females (Table S2; OR = 2.30, p < 0.01; Fig. 3A), and reproductive bats having a marginally lower likelihood of infection than nonreproductive bats (Table S2; OR = 0.55, p = 0.05; Fig. 3C). There was no significant effect of site or age class (Table S2). The fixed effects accounted for 14% of the variance in infection status (R^2^_m_= 0.14) while bat ID accounted for 7% (R^2^ = 0.21), indicating that recaptured bats (n = 59) had little effect on the model predictions.

**Figure 3.**
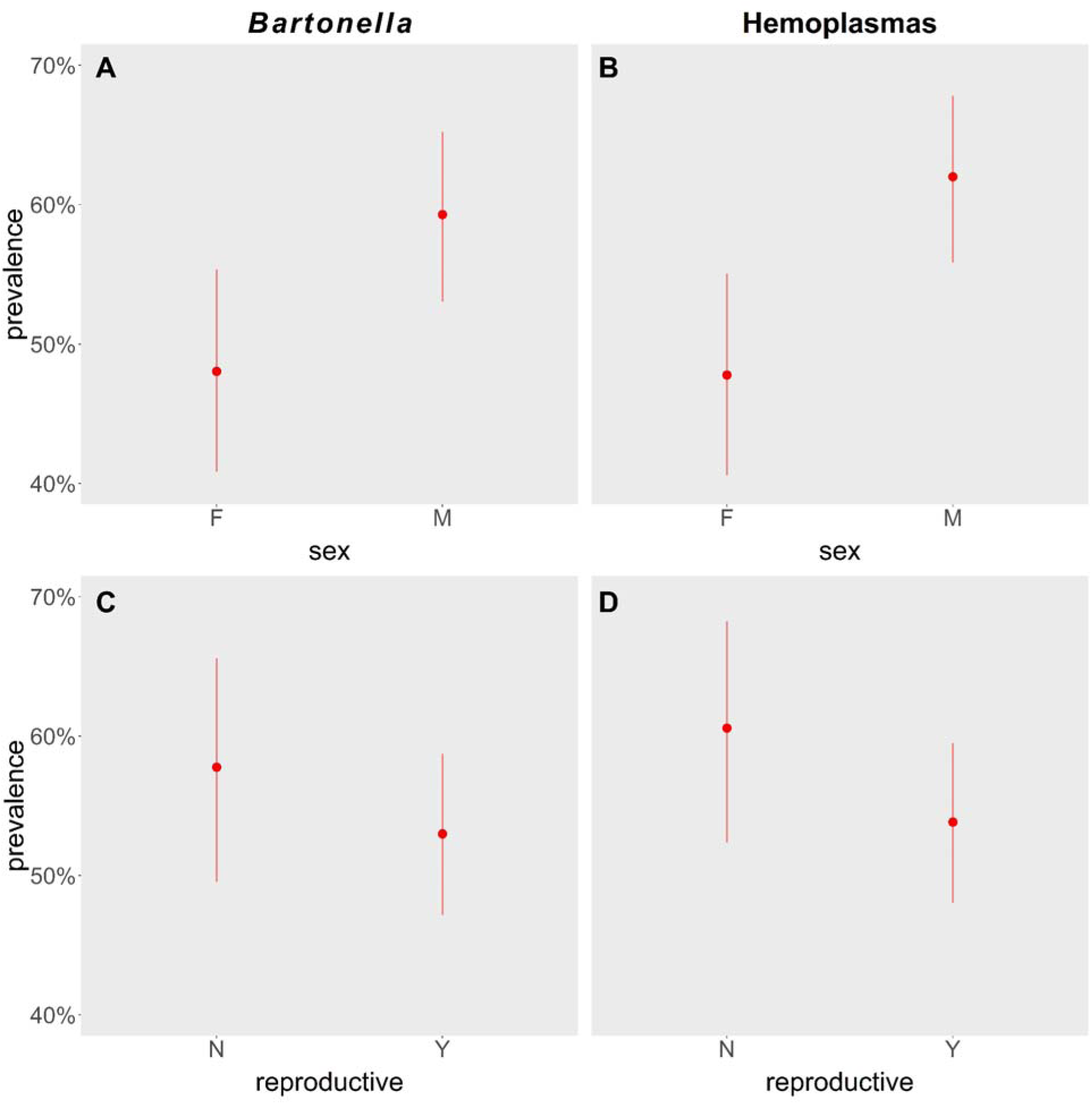
*Bartonella* (A–B) and hemoplasma (C–D) prevalence by sex and reproductive status. Point estimates are shown with 95% confidence intervals (Wilson interval).

**Table 1.**
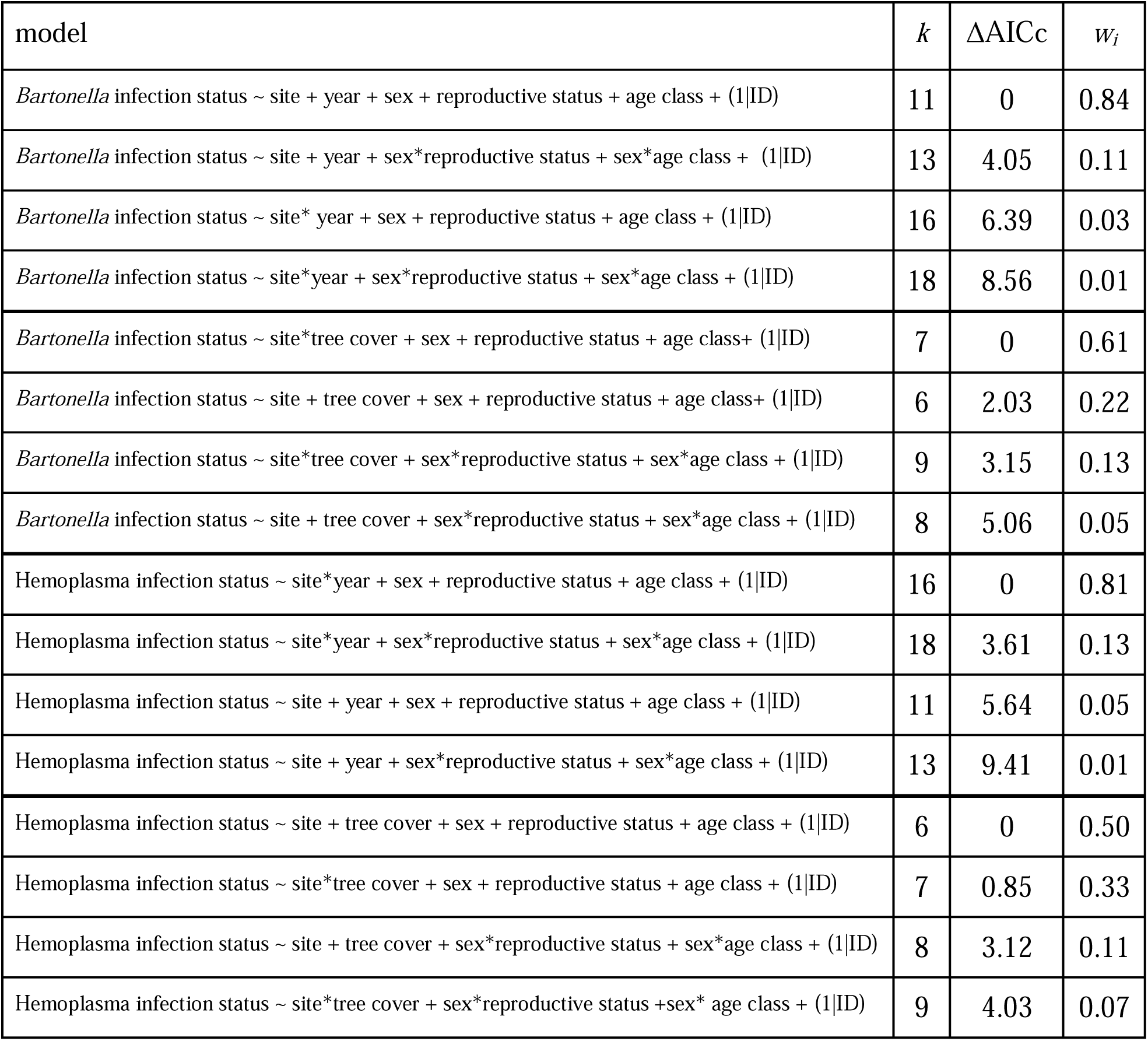
Competing suites of spatiotemporal and tree cover GLMMs for *Bartonella* and hemoplasma infection status. Within each suite, models are ranked by ΔAICc with the number of coefficients (*k*) and Akaike weights (*w_i_*).

| model | $k$ | $\Delta\text{AICc}$ | $w_i$ |
| --- | --- | --- | --- |
| *Bartonella* infection status ~ site + year + sex + reproductive status + age class + (1\|ID) | 11 | 0 | 0.84 |
| *Bartonella* infection status ~ site + year + sex*reproductive status + sex*age class + (1\|ID) | 13 | 4.05 | 0.11 |
| *Bartonella* infection status ~ site* year + sex + reproductive status + age class + (1\|ID) | 16 | 6.39 | 0.03 |
| *Bartonella* infection status ~ site*year + sex*reproductive status + sex*age class + (1\|ID) | 18 | 8.56 | 0.01 |
| *Bartonella* infection status ~ site*tree cover + sex + reproductive status + age class+ (1\|ID) | 7 | 0 | 0.61 |
| *Bartonella* infection status ~ site + tree cover + sex + reproductive status + age class+ (1\|ID) | 6 | 2.03 | 0.22 |
| *Bartonella* infection status ~ site*tree cover + sex*reproductive status + sex*age class + (1\|ID) | 9 | 3.15 | 0.13 |
| *Bartonella* infection status ~ site + tree cover + sex*reproductive status + sex*age class + (1\|ID) | 8 | 5.06 | 0.05 |
| Hemoplasma infection status ~ site*year + sex + reproductive status + age class + (1\|ID) | 16 | 0 | 0.81 |
| Hemoplasma infection status ~ site*year + sex*reproductive status + sex*age class + (1\|ID) | 18 | 3.61 | 0.13 |
| Hemoplasma infection status ~ site + year + sex + reproductive status + age class + (1\|ID) | 11 | 5.64 | 0.05 |
| Hemoplasma infection status ~ site + year + sex*reproductive status + sex*age class + (1\|ID) | 13 | 9.41 | 0.01 |
| Hemoplasma infection status ~ site + tree cover + sex + reproductive status + age class + (1\|ID) | 6 | 0 | 0.50 |
| Hemoplasma infection status ~ site*tree cover + sex + reproductive status + age class + (1\|ID) | 7 | 0.85 | 0.33 |
| Hemoplasma infection status ~ site + tree cover + sex*reproductive status + sex*age class + (1\|ID) | 8 | 3.12 | 0.11 |
| Hemoplasma infection status ~ site*tree cover + sex*reproductive status +sex* age class + (1\|ID) | 9 | 4.03 | 0.07 |

### Effect of tree cover on Bartonella infection

For the subset of our *Bartonella* data that included tree cover (2017–2022), the GLMM with an interaction between site and the area of tree cover in the matrix and additive effects of sex, reproductive status, and age class was superior (Table 1). Infection was best predicted by the interaction between site and tree cover, and by sex with reproductive status as a marginal predictor (Table S3). While the area of tree cover in the matrix had no effect on *Bartonella* infection in KK (= 0.02, CI =-0.47 to 0.51), we found that decreased tree cover was associated with a higher infection likelihood in the LAR (= 0.60, CI =-0.95 to-0.25) (Fig. 4). As in the GLMMs fit to the full dataset, males were more likely to be infected by females (OR = 2.38, p < 0.01), reproductive bats were marginally less likely to have *Bartonella* infections than nonreproductive bats (OR = 0.60, p = 0.08) and there was no effect of age class (Table S3). The fixed effects accounted for 11% of the variance in the data (R^2^_m_= 0.11), with no additional contribution from bat ID (R^2^_c_= 0.11).

**Figure 4.**
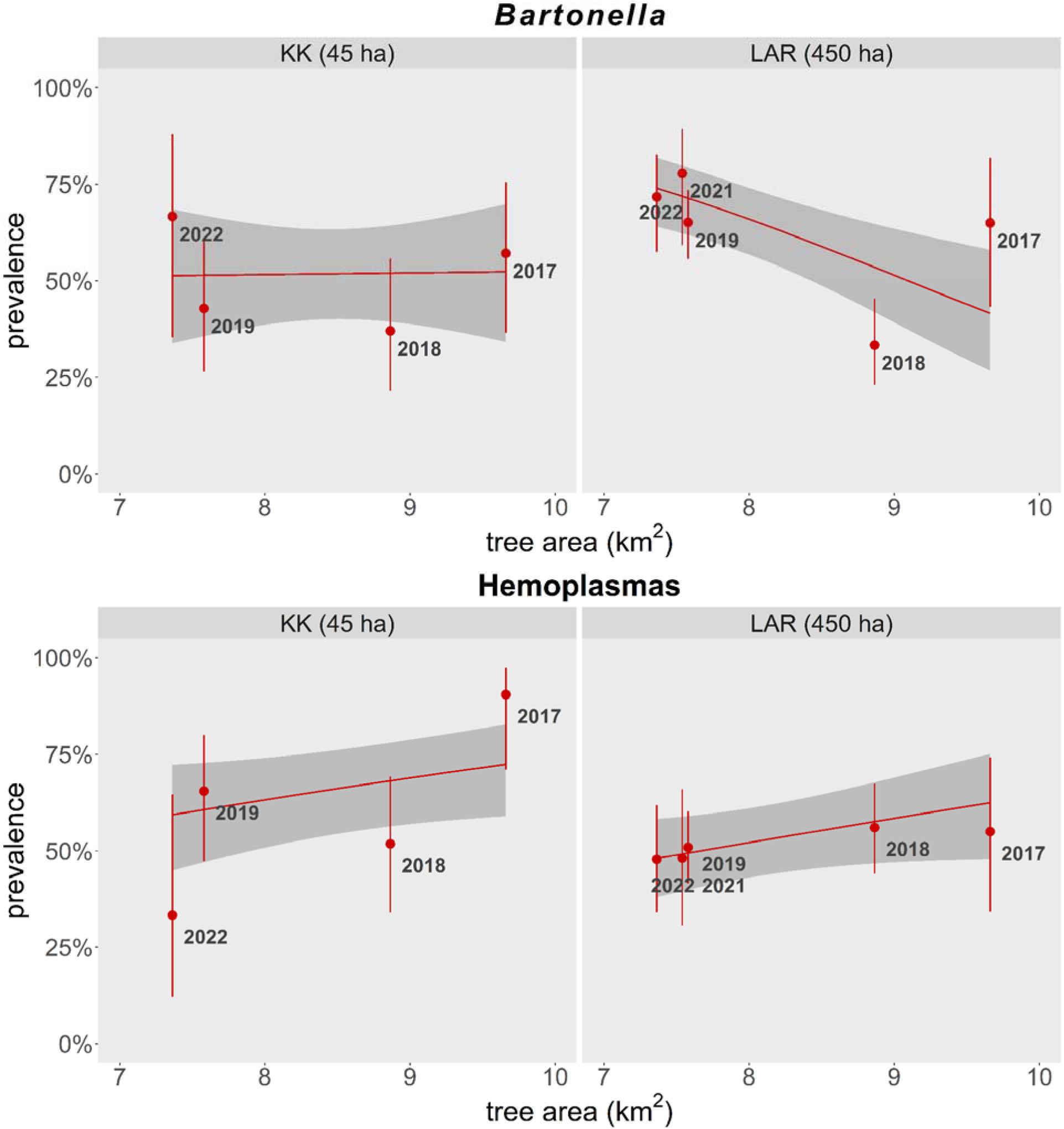
*Bartonella* and hemoplasma infection prevalence by tree cover in each site from 2017-2022.

### Bartonella genotype diversity (2015–2022)

We identified 16 different *Bartonella* genotypes in this study (Fig. S2). Nine of these (DR1, 2, 4– 6,8–11) had been identified in our previous study of vampire bats (Becker et al., 2018a), and we had previously identified one genotype (TB1) in Mexican free-tailed bats (*Tadarida brasiliensis)* in Oklahoma (Becker et al., 2025). The remaining six genotypes (DR12–17) were novel. The genotypes identified were highly similar to reference sequences from bat and bat fly species, but not reference sequences isolated from humans, domestic animals, or other wildlife. The two sequences assigned to DR12 (GenBank accessions PQ758896 and PQ758863) were nearly identical (>99%) to sequences previously found in streblid bat fly (*Megistopoda proxima*) found on *Sturnira parvidens* from Costa Rica (Genbank accession MH234363). Meanwhile, five sequences (GenBank accessions PQ758813, PQ758901, PQ758909, PQ758910, PQ758950) were assigned to DR13 and were 99–100% similar to other bartonellae previously found in vampire bats from Brazil (GenBank accession PP715670), Mexico (GenBank accession MF467787), and Guatemala (GenBank accession MN529504). Only one sequence each made up genotypes DR14, DR15, DR16, and DR17. The DR14 sequence (GenBank accession PQ758852) was 96.2% similar to a *Bartonella* sp. from *Myotis nigricans* from Guatemala (GenBank accession MN529480). DR15 (GenBank accession PQ758988) was only 88.1% similar to *Bartonella* sp. from a bat fly in China (GenBank accession MZ208693). DR16 (GenBank accession PQ758964) was 93.7% similar to a *Bartonella* sp. from *Carollia perspicillata* from Costa Rica (GenBank accession MH234350). Lastly, DR17 (GenBank accession PQ758949) was 93.1% similar to a *Bartonella* sp. from *Glossophaga soricina* from Costa Rica (GenBank accession MH234356).

As most of the genotypes were observed <10 times each across the entire study period, only those with at least 10 observations were included in subsequent analyses: DR1 (n = 25), DR2 (n = 21), DR8 (n = 32), DR9 (n = 31), and DR11 (n = 56), all of which were identified previously (Becker et al., 2018a). The frequencies of DR8 (χ*^2^* = 17.4, p < 0.01), DR9 (χ*^2^* = 26.1, p < 0.01), and DR11 (χ*^2^* = 56.5, p < 0.01) all varied significantly across years, but those of DR1 χ = 7.40, p = 0.19) and DR2 (χ = 6.71, p = 0.24) did not. These five dominant genotypes oscillated through periods of equilibrium and periods of dominance by one or more genotypes. The most recent years of sampling saw no significant differences in genotype frequency during 2021 (χ^2^ = 4.47, p = 0.35) or 2022 (χ^2^ = 1.67, p = 0.80). The same was true for 2015 (χ*^2^* = 0.67, p = 0.96) and 2017 (χ*^2^* = 4.00 p = 0.41), with periods of significant differences in *Bartonella* genotype frequencies in 2018 (χ*^2^* = 12.4, p = 0.01) and 2019 (χ*^2^* = 24.5 p < 0.01). When DR11 was 22% and 15% more prevalent than the next most prevalent genotype, respectively (Fig. S3). Only DR8 was detected in 2016, preventing statistical comparison but demonstrating a period of dominance by DR8 over others.

### Spatiotemporal predictors of Bartonella genotypes

The model that best predicted infection with each respective genotype among *Bartonella*-infected bats included the additive effects of site, year, sex, reproductive, status, and age class (Table 2). Significant predictors were not identified for DR2 or DR11, but age class had a marginal effect on DR1 and DR9 infections (Table S4) in which adults were marginally less likely to be infected with DR1 (OR = 2.86; p = 0.08), but marginally more likely to be infected with DR9 (OR = 0.26, p = 0.06) than subadults (Fig. S4A-B). Sex was a marginal predictor for DR8 infection (Table S4); females were marginally less likely to be infected by DR8 than males (OR = 2.62, p = 0.08; Fig. S4C). The fixed effects accounted for 55–57% of the variance in the data for DR1 (R^2^_m_= 0.56), DR2 (R^2^_m_= 0.54), DR9 (R^2^_m_= 0.57), and DR11 (R^2^_m_= 0.55) respectively, and 10% of the data for DR8 (R^2^_m_= 0.10). No additional variance was attributed to bat ID (DR1 R^2^ = 0.56; DR2 R^2^ = 0.54; DR8 R^2^ = 0.10; DR9 R^2^ = 0.57; DR11 R^2^ = 0.55).

**Table 2.** Competing suites of spatiotemporal GLMMs for infection status per each *Bartonella* genotype. Within each suite, models are ranked by ΔAICc with the number of coefficients (*k*) and Akaike weights (*w_i_*).

| model | $k$ | $\Delta\text{AICc}$ | $w_i$ |
| --- | --- | --- | --- |
| DR1 infection status ~ site + year + sex + reproductive status + age class + (1\|ID) | 11 | 0 | 0.81 |
| DR1 infection status ~ site*year + sex + reproductive status + age class + (1\|ID) | 16 | 4.20 | 0.10 |
| DR1 infection status ~ site + year + sex* reproductive status + sex*age class + (1\|ID) | 13 | 4.61 | 0.08 |
| DR1 infection status ~ site*year + sex*reproductive status + sex*age class + (1\|ID) | 18 | 8.87 | 0.01 |
| DR2 infection status ~ site + year + sex + reproductive status + age class + (1\|ID) | 11 | 0 | 0.79 |
| DR2 infection status ~ site + year + sex* reproductive status + sex* age class + (1\|ID) | 13 | 3.96 | 0.11 |
| DR2 infection status ~ site*year + sex + reproductive status + age class + (1\|ID) | 16 | 4.40 | 0.09 |
| DR2 infection status ~ site*year + sex*reproductive status + sex*age class + (1\|ID) | 18 | 8.78 | 0.01 |
| DR8 infection status ~ site + year + sex*reproductive status + sex*age class + (1\|ID) | 13 | 0 | 0.53 |
| DR8 infection status ~ site + year + sex + reproductive status + age class + (1\|ID) | 11 | 0.65 | 0.39 |
| DR8 infection status ~ site*year + sex*reproductive status + sex*age class + (1\|ID) | 18 | 5.15 | 0.04 |
| DR8 infection status ~ site*year + sex + reproductive status + age class + (1\|ID) | 16 | 5.27 | 0.04 |
| DR9 infection status ~ site + year + sex + reproductive status + age class + (1\|ID) | 11 | 0 | 0.60 |
| DR9 infection status ~ site + year + sex* reproductive status + sex* age class + (1\|ID) | 13 | 1.12 | 0.34 |
| DR9 infection status ~ site*year + sex + reproductive status + age class + (1\|ID) | 16 | 5.42 | 0.04 |
| DR9 infection status ~ site*year + sex*reproductive status + sex*age class + (1\|ID) | 18 | 6.98 | 0.02 |
| DR11 infection status ~ site + year + sex + reproductive status + age class + (1\|ID) | 11 | 0 | 0.76 |
| DR11 infection status ~ site + year + sex*reproductive status + sex*age class + (1\|ID) | 13 | 3.10 | 0.16 |
| DR11 infection status ~ site*year + sex + reproductive status + age class + (1\|ID) | 16 | 4.71 | 0.07 |
| DR11 infection status ~ site*year + sex*reproductive status + sex*age class + (1\|ID) | 18 | 8.09 | 0.01 |

### Effect of tree cover on Bartonella genotypes

The model that best predicted infections with DR1, DR2, DR8, and DR11 among *Bartonella*-infected bats included the additive effects of site, tree area, sex, reproductive status, and age class (Table 3). The area of tree cover in the matrix was the only significant predictor of DR1 infection (Table S5) in which decreasing tree cover was associated with an decreased likelihood of infection (OR = 0.49, p = 0.01) (Fig. 5). We did not identify significant predictors of DR2 or DR11 infection, but sex marginally predicted DR8 infection status (Table S5). Males were marginally more likely to be infected with DR8 than females (OR = 2.63, p = 0.08). The model that best predicted DR9 infections included the interaction between site and tree area as well as additive effects of sex, reproductive status, and age class (Table 3). The interaction between site and tree cover was a significant predictor of DR9 infection (Table S5) and age class was a marginal predictor. While subadults were less likely to be infected with DR9 compared to adults (OR = 0.26, p = 0.06), we did not identify post-hoc differences in infection likelihood between sites by tree cover (KK = 1.03, CI =-0.28 to 2.33; LAR =-0.59, CI =-1.41 to 0.24; Fig. 5). The fixed effects included in the models accounted for 10%, 3%, 9%, 15%, and 3% of the variance in the data for DR1 (R^2^_m_= 0.10), DR2 (R^2^_m_= 0.03), DR8 (R^2^_m_= 0.09), DR9, (R^2^_m_= 0.15) and DR11 (R^2^_m_= 0.03), respectively. No additional variance was attributed to bat ID for DR1 (R^2^_c_= 0.10), DR2 (R^2^_c_= 0.03), DR8 (R^2^_c_= 0.09), DR9 (R^2^_c_= 0.15), or DR11 (R^2^_c_= 0.03).

**Figure 5.**
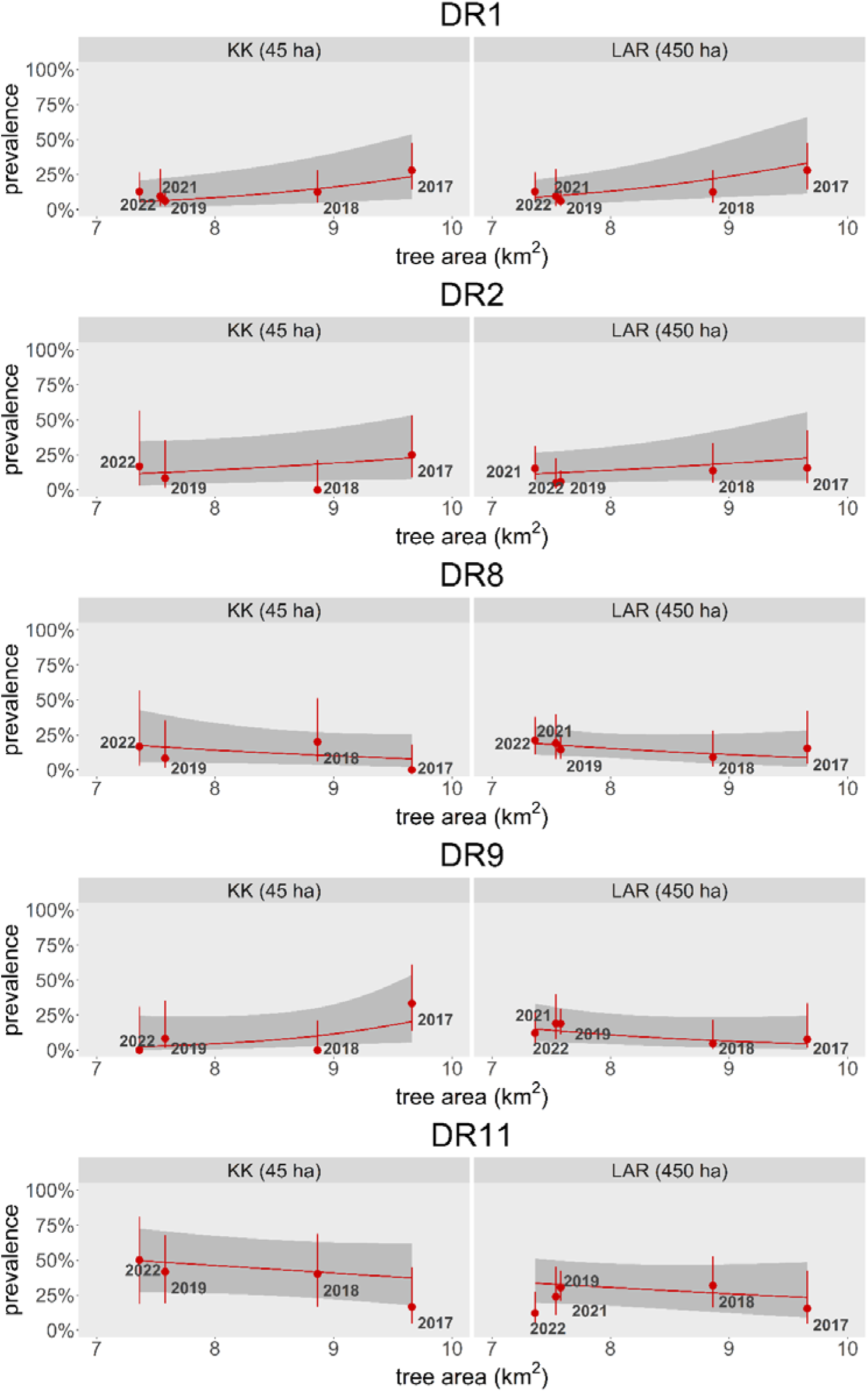
Prevalence of *Bartonella* genotype infections by tree cover from 2017-2022.

**Table 3.**
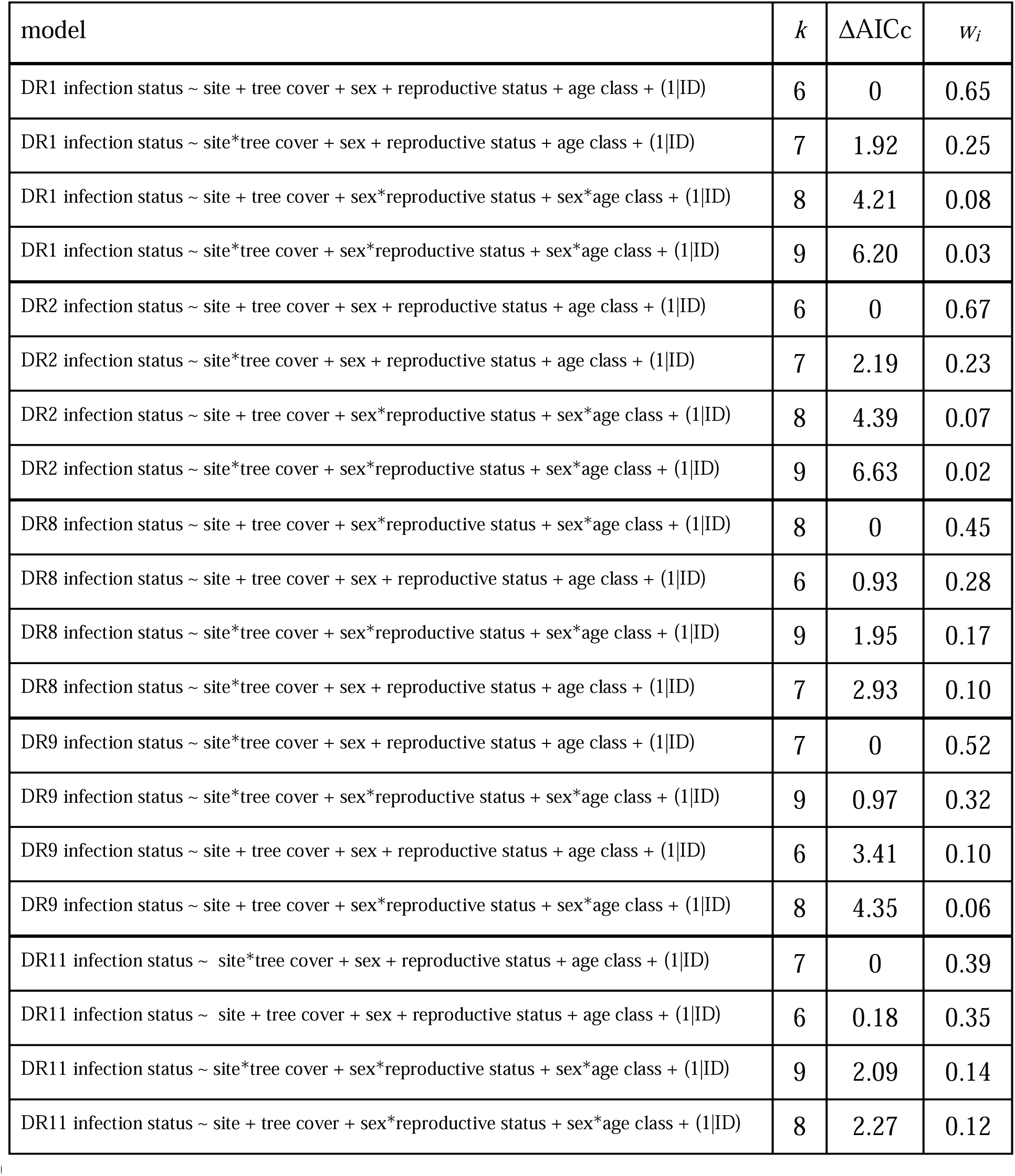
Competing suites of tree cover GLMMs for infection status per each *Bartonella* genotype. Within each suite, models are ranked by ΔAICc with the number of coefficients (*k*) and Akaike weights (*w_i_*).

### Bartonella infection status switch and genotype switching among recaptured bats

Across the entire study period, 59 bats were recaptured at least once (145 total captures). Of the recaptured bats, 19 were never found to be infected with *Bartonella*. Twenty-six bats switched infection status during our study, 13 individuals switched from positive to negative, seven switched from negative to positive, four switched from positive to negative and then back to positive, and two switched from negative to positive and then back to negative. Eleven bats with multiple years of *Bartonella* infections switched genotypes, which could indicate loss and gain of infection or altered within-host dynamics that allowed a less-abundant genotype to dominate.

While two different genotypes were detected sequentially in 10 of the 40 recaptured bats with *Bartonella* infections, only one recaptured bat harbored infections with three different genotypes across the study period, and only one genotype was detected in 29 of the recaptured bats (Fig. S5).

Bats with more capture events were more likely to switch between *Bartonella* genotypes (β = 1.98, p < 0.01). Additionally, bats with a higher number of capture events were also more likely to host a higher number of *Bartonella* genotypes across the study period (β = 0.54, p = 0.02) (Fig. S6). We did not identify any predictors of *Bartonella* infection status switching (Table S6).

### Spatiotemporal predictors of hemoplasma infection

Among four GLMMs predicting hemoplasma infection status across all years (2015–2022), the most competitive model included the interaction between site and year as well as additive effects of sex, reproductive status, and age class (Table 1). Infection was best predicted by sex, the interaction between site and year, and reproductive status (Table S2). Hemoplasma infections were marginally more likely in KK in 2017 compared to the LAR in 2019 (OR = 10.5, p = 0.1), 2021 (OR = 11.7, p = 0.1), and 2022 (OR = 11.4, p = 0.1) and compared to KK in 2015 (OR = 0.07, p = 0.1), 2016 (OR = 0.06, p = 0.1), and 2022 (OR = 24.2, p = 0.1) (Fig. 2B). Males were more likely to be infected than females (OR = 2.38, p < 0.01; Fig 3B), and non-reproductive bats were more likely to be infected than reproductive bats (OR = 0.54, p = 0.04; Fig. 3D). There was no effect of age class on hemoplasma infection (Table S2). The fixed effects accounted for 15% of the variance in infection status (R^2^_m_ = 0.15), with bat ID explaining an additional 9% of the variance (R^2^ = 0.24).

### Effect of tree cover on hemoplasma infection

For the subset of our hemoplasma data that included tree cover (2017–2022), the GLMM with additive effects of site, tree cover, sex, reproductive status, and age class was superior (Table 1). Infection was best predicted by sex and reproduction status, while site and tree cover were marginal predictors (Table S3). As in our analyses with the full dataset, males were more likely to be infected than females (OR = 2.02, p = <0.01), while non-reproductive bats had higher likelihoods of infection compared to reproductive bats (OR = 0.47, p = <0.01). Hemoplasma prevalence was marginally higher in KK than in the LAR (OR = 0.63, p = 0.1), and decreased tree cover was marginally associated with decreased likelihood of infection (OR = 0.77, p = 0.08) (Fig. 4). There was no significant effect of age class on infection status (Table S3). The fixed effects included in the model accounted for 8% of the variance in infection status (R^2^_m_= 0.08). No additional variance was attributed to bat ID (R^2^_c_= 0.08).

### Hemoplasma genotypic diversity (2015–2022)

As suggested by our earlier work (Volokhov et al., 2017; Becker et al., 2020b; DeAnglis et al., 2024), three genotypes (VB1, VB2, VB3) comprised the majority of hemoplasma infections (240 of 242 total hemoplasma-positive bats). In addition, one bat (GenBank accession OQ385174) was infected with a potentially novel hemoplasma genotype with a 16S rRNA gene sequence 99% similar to that of genotype MR1 (GenBank accession MH245174), which was previously found in *Molossus nigricans* and *Molossus alvarezi* (Becker et al., 2020b; Volokhov et al., 2023). Another (GenBank accession OQ546513) was identified as belonging to genotype EF1 (GenBank accession MH245131), which typically infects *Neoeptesicus furinalis,* but has also been found in *Glossophage soricina* and *Saccopteryx bilineata* (Becker et al., 2020b; Volokhov et al., 2023; Fig. S7). As previously reported, a third 16S rRNA sequence (GenBank accession KY932724) was 98% similar to *Mycoplasma moatsii* (GenBank accession NR_025186), a non-hemotropic species, and therefore not considered to be a hemoplasma (Volokhov et al., 2017).

These genotypes did not share a high degree of genetic similarity to reference sequences from humans, domestic animals, other wildlife, or potential arthropod vectors (Fig. S7). Across all bats in our study, 17 were coinfected with multiple hemoplasma genotypes and were classified according to the dominant genotype. Only infections belonging to the dominant genotypes were included in subsequent analyses: VB1 (n = 106), VB2 (n = 85), and VB3 (n = 49). Amplification of paired partial 23S rRNA and/or *rpoB* genes for samples belonging to these 16S rRNA genotypes (see Table S1) suggested at least two novel *Candidatus* hemoplasma species circulate in vampire bats. Based on 99.9% identity of three 23S rRNA sequences (OQ518936, OQ518940, OQ518941) and 99.9–100% identity of paired 16S rRNA sequences included in the VB1 genotype (MH245176, OQ546521, OQ546522), first detected in the LAR, we propose the name *Candidatus* Mycoplasma haematolamanidesmodi sp. nov.; paired *rpoB* sequences in two of these three bats also showed 100% identity (OQ554324, OQ554325), further confirming this proposed hemoplasma species. Similarly, given 99.9–100% identity among three 23S rRNA sequences (OQ518937, OQ518938, OQ518946) and 99.1–99.4% identity of paired 16S rRNA sequences included in the VB2 genotype (OQ385162, OQ546569, OQ546517), first detected in KK, we propose the name *Candidatus* Mycoplasma haematokakabidesmodi sp. nov.

The frequency of all three of the main hemoplasma genotypes varied significantly among years (VB1 χ*^2^* = 22.2, p < 0.01; VB2 χ*^2^* = 64.5, p < 0.01; VB3 χ*^2^* = 26, p < 0.01). The frequency of VB1 decreased from 2017 to 2019 while that of VB2 increased during the same timespan. The frequency of VB3 has gradually increased throughout the study period, and the three genotypes have reached an equilibrium in the most recent years of sampling, with no significant differences in frequency during 2021 (χ^2^ = 1.08, p = 0.58) or 2022 (χ^2^ = 0.75, p = 0.69). The only other year without significant differences in hemoplasma genotype frequencies was 2018 (χ*^2^*= 0.82, p = 0.66). While genotype frequencies were marginally different in 2015 (χ*^2^* = 5.76, p = 0.06), they varied significantly during 2016 (χ*^2^* = 22.6, p < 0.01), 2017 (χ*^2^*= 13.6, p < 0.01), and 2019 (χ*^2^* = 10.1, p < 0.01) (Fig. S8).

### Spatiotemporal predictors of hemoplasma genotypes

Among the four models predicting infection with each respective hemoplasma genotype among infected bats across all years (2015–2022), the GLMM that best predicted infection across genotypes included the additive effects of site, year, sex, reproductive status, and age class (Table 4). VB1 infection was best predicted by year, while age class was a marginal predictor (Table S7). VB1 infection was significantly more likely in 2016 compared to 2018 (OR = 6.71, p < 0.01), 2019 (OR = 6.18, p < 0.01), and 2022 (OR = 4.65, p = 0.04) and significantly more likely in 2017 compared to 2018 (OR = 5.07, p = 0.01) and 2019 (OR = 4.67, p = 0.01) (Fig. S8). Adults with hemoplasma infections were marginally less likely to carry VB1 than infected subadults (OR = 2.17, p = 0.05; Fig. S9A). There were no effects of site, sex, or reproductive status on VB1 infection (Table S7). The fixed effects accounted for 16% of the variance in the data (R^2^_m_= 0.16), while no additional variance was attributed to bat ID (R^2^ = 0.16).

**Table 4.**
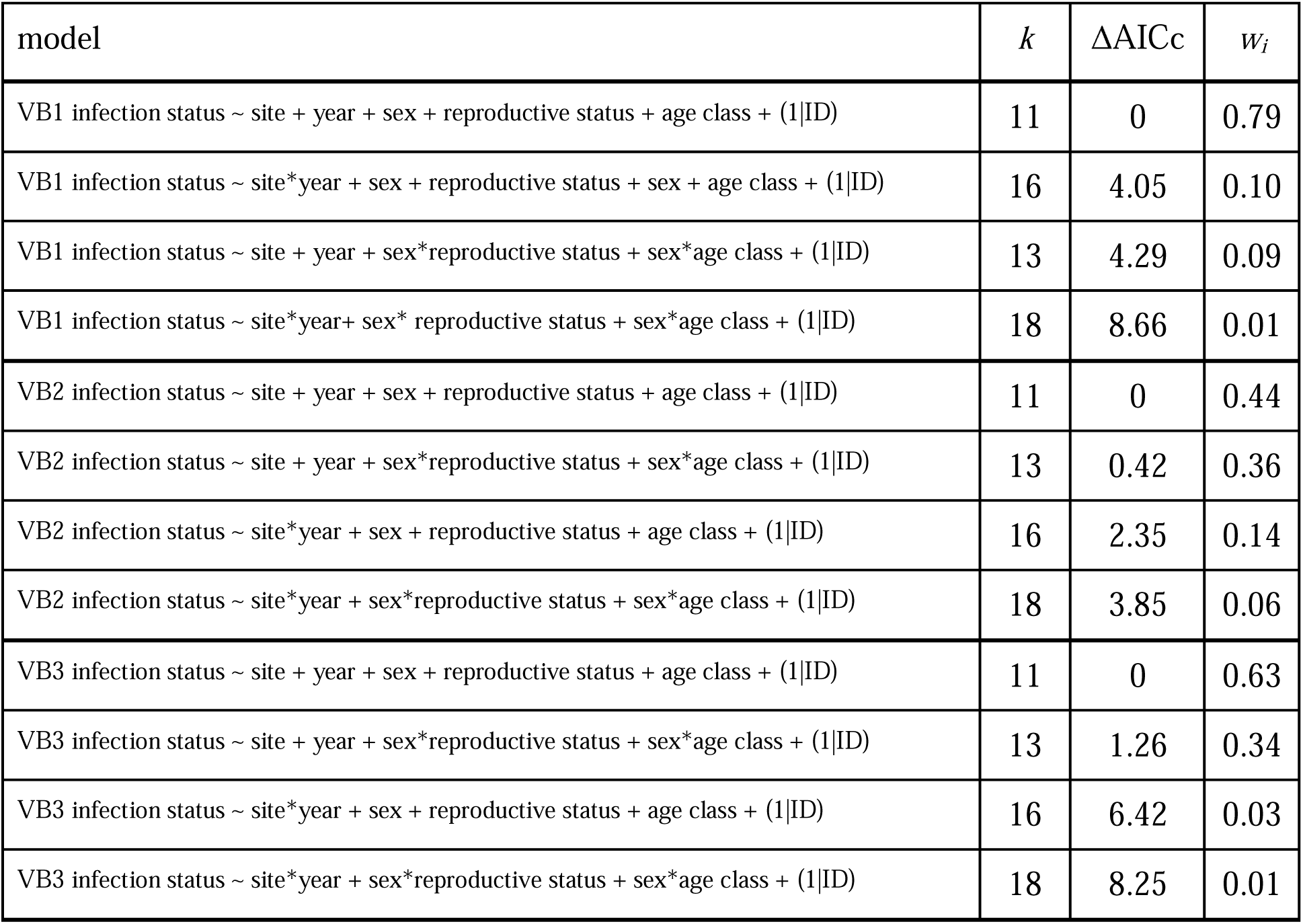
Competing suites of spatiotemporal GLMMs for infection status per each hemoplasma genotype. Within each suite, models are ranked by ΔAICc with the number of coefficients (*k*) and Akaike weights (*w_i_*).

| model | $k$ | $\Delta\text{AICc}$ | $w_i$ |
| --- | --- | --- | --- |
| VB1 infection status ~ site + year + sex + reproductive status + age class + (1\|ID) | 11 | 0 | 0.79 |
| VB1 infection status ~ site*year + sex + reproductive status + sex + age class + (1\|ID) | 16 | 4.05 | 0.10 |
| VB1 infection status ~ site + year + sex*reproductive status + sex*age class + (1\|ID) | 13 | 4.29 | 0.09 |
| VB1 infection status ~ site*year+ sex* reproductive status + sex*age class + (1\|ID) | 18 | 8.66 | 0.01 |
| VB2 infection status ~ site + year + sex + reproductive status + age class + (1\|ID) | 11 | 0 | 0.44 |
| VB2 infection status ~ site + year + sex*reproductive status + sex*age class + (1\|ID) | 13 | 0.42 | 0.36 |
| VB2 infection status ~ site*year + sex + reproductive status + age class + (1\|ID) | 16 | 2.35 | 0.14 |
| VB2 infection status ~ site*year + sex*reproductive status + sex*age class + (1\|ID) | 18 | 3.85 | 0.06 |
| VB3 infection status ~ site + year + sex + reproductive status + age class + (1\|ID) | 11 | 0 | 0.63 |
| VB3 infection status ~ site + year + sex*reproductive status + sex*age class + (1\|ID) | 13 | 1.26 | 0.34 |
| VB3 infection status ~ site*year + sex + reproductive status + age class + (1\|ID) | 16 | 6.42 | 0.03 |
| VB3 infection status ~ site*year + sex*reproductive status + sex*age class + (1\|ID) | 18 | 8.25 | 0.01 |

Infection with VB2 among hemoplasma-infected bats was best predicted by year, while the effects of reproductive status was a marginal predictor (Table S7). VB2 infection was marginally less likely in 2016 (OR = 0.23, p = 0.05) and significantly less likely in 2017 (OR = 0.16, p = 0.03) compared to 2019. Additionally, VB2 infection was marginally more likely in 2015 compared to 2017 (OR = 5.05, p = 0.1), marginally less likely in 2017 compared to 2018 (OR = 0.23, p = 0.08), and marginally more likely in 2019 compared to 2022 (OR = 4.15, p = 0.07) (Fig. S8). Reproductive (OR = 2.09, p = 0.06) bats were marginally more likely to be infected with VB2 than nonreproductive bats (Fig. S9B). There were no effects of site, sex, or age class (Table S7). The fixed effects accounted for 18% of the variance in the data (R^2^_m_= 0.18). No additional variance was attributed to bat ID (R^2^ = 0.18).

Year was a marginal predictor of VB3 infection likelihood, but we did not identify post hoc differences between years nor did we identify other predictors (Table S7). Fixed effects accounted for only 16% of variance (R^2^_m_= 0.17). Bat ID did not explain further variance (R^2^ = 0.17).

### Effect of tree cover on hemoplasma genotypes

For the subset of hemoplasma*-*infected bats for which tree cover data were available, (2017– 2022), the model that best predicted infection with VB1 included the additive effects of site, tree cover, sex, reproductive status, and age class (Table 5).VB1 infection was best predicted by tree cover (Table S8), where decreasing tree cover had a negative effect on infection likelihood (OR = 0.66, p = 0.03) (Fig. 6). While there was no effect of site, sex, or reproductive status on the odds of VB1 infection, there was a marginal effect of age class (Table S8). As in the spatiotemporal model, adults were marginally less likely to be infected than subadults (OR = 2.06, p = 0.09) The fixed effects accounted for 5% of the variance in the data (R^2^_m_= 0.05), with no additional variance attributed to bat ID (R^2^ = 0.05).

**Figure 6.**
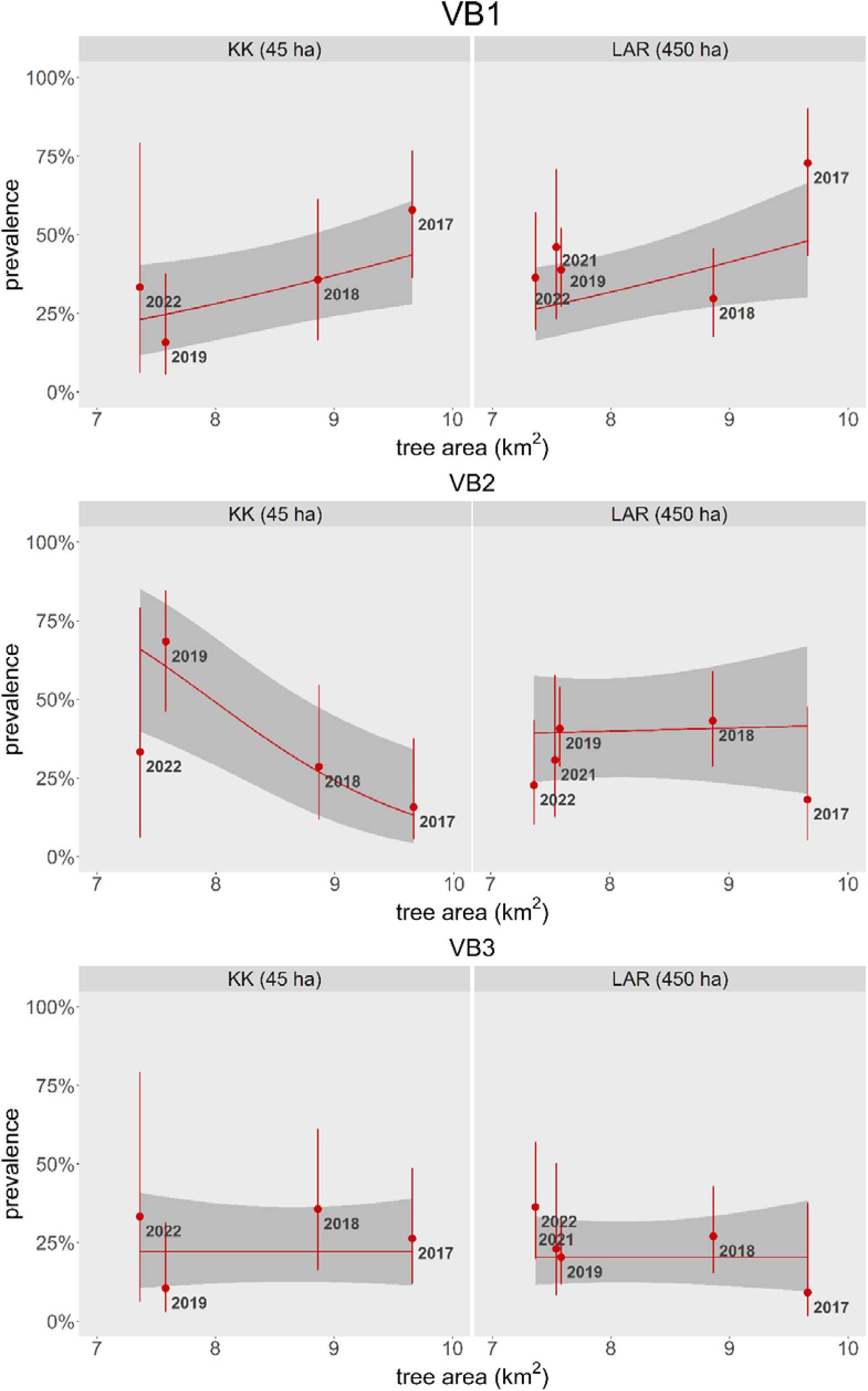
Prevalence of hemoplasma genotype infections by tree cover from 2017-2022. Point estimates are shown with 95% confidence intervals (Wilson’s interval).

**Table 5.** Competing suites of tree cover GLMMs for infection status per each hemoplasma genotype. Within each suite, models are ranked by ΔAICc with the number of coefficients (*k*) and Akaike weights (*w_i_*).

| model | $k$ | $\Delta\text{AICc}$ | $w_i$ |
| --- | --- | --- | --- |
| VB1 infection status ~ site*tree cover + sex + reproductive status + age class + (1\|ID) | 7 | 0 | 0.56 |
| VB1 infection status ~ site + tree cover + sex + reproductive status + age class + (1\|ID) | 6 | 0.99 | 0.34 |
| VB1 infection status ~ site*tree cover + sex*reproductive status + sex*age class + (1\|ID) | 9 | 4.44 | 0.06 |
| VB1 infection status ~ site + tree cover + sex*reproductive status + sex*age class + (1\|ID) | 8 | 5.37 | 0.04 |
| VB2 infection status ~ site*tree cover + sex + reproductive status + age class + (1\|ID) | 7 | 0 | 0.65 |
| VB2 infection status ~ site*tree cover + sex*reproductive status + sex*age class + (1\|ID) | 9 | 1.62 | 0.29 |
| VB2 infection status ~ site + tree cover + sex + reproductive status + age class + (1\|ID) | 6 | 5.50 | 0.04 |
| VB2 infection status ~ site + tree cover + sex*reproductive status + sex*age class + (1\|ID) | 8 | 6.81 | 0.02 |
| VB3 infection status ~ site + tree cover + sex + reproductive status + age class + (1\|ID) | 6 | 0 | 0.37 |
| VB3 infection status ~ site*tree cover + sex + reproductive status + age class + (1\|ID) | 7 | 0.41 | 0.30 |
| VB3 infection status ~ site + tree cover + sex*rep + sex*age class + (1\|ID) | 8 | 1.42 | 0.18 |
| VB3 infection status ~ site*tree cover + sex*reproductive status + sex*age class + (1\|ID) | 9 | 1.92 | 0.14 |

The model that best predicted infection with VB2 included the interaction between site and tree area along with sex, reproductive status, and age class (Table 5). VB2 infection was best predicted by the interaction between tree area and site (Table S8). Decreasing tree cover positively affected VB2 infection in KK (= 1.11; CI =-1.80 to-0.41) but not in the LAR (= 0.04; CI =-0.43 to 0.51) (Fig. 6). We detected no significant effect of sex, reproductive status, or age class (Table S8). The fixed effects accounted for 12% of the variance in infection status (R^2^_m_= 0.12). No additional variance was attributed to bat ID (R^2^ = 0.12).

The top model for VB3 among hemoplasma-infected bats included additive effects of site, tree cover, sex, reproductive status, and age class (Table 5). However, we identified no significant predictors of VB3 infection (Table S8). The fixed effects explained 3% of the variance in the data (R^2^_m_= 0.03), with no additional contribution from bat ID (R^2^ = 0.03).

### Hemoplasma infection status switching and genotype switching among recaptured bats

We detected hemoplasma infections from 48 of the 59 bats recaptured during this study. Twenty-five bats changed infection statuses, with 14 switching from positive to negative, eight switching from negative to positive, two switching from positive to negative and then back to positive, and one switching from negative to positive and then back to negative. Twelve bats with multiple hemoplasma infections switched between genotypes. Most recaptured bats (n = 29, 60%) only had one detected genotype during the study period. Twelve bats were infected with two hemoplasma genotypes at some point during the study period, and none carried more than two genotypes (Fig. S10). However, we did not identify any predictors of genotype switching, infection status switching, or the number of genotypes identified in a recaptured bat across the study period (Table S9).

## Discussion

Habitat fragmentation typically has detrimental effects on wildlife, including increased risk of pathogen exposure (Haddad et al., 2015, Gibb et al, 2024). However, testing for such effects is difficult, as the effects of habitat fragmentation can take years to manifest (Heckley et al., 2023) and are challenging to distinguish from effects of habitat loss (Laurance, 2008; Didham et al., 2012). Here, we show that despite similarities in prevalence within our studied vampire bat populations, bartonellae and hemoplasmas varied in their spatiotemporal dynamics within the changing landscape and in their responses to forest loss within the matrix surrounding two habitat fragments. Additionally, responses to the fixed effects included in our models varied by genotype within both of the two respective pathogens, suggesting that the effects of processes such as habitat fragmentation on infectious disease dynamics may not be easily generalizable across or within pathogens.

### Bartonellae and hemoplasmas differ in their spatiotemporal dynamics

As expected, we observed differing infection dynamics between bartonellae and hemoplasmas, which may be due to differences in the relative evidence supporting vector-borne and direct transmission (Willi et al., 2007; Judson et al., 2015; Becker et al., 2018a). Vampire bats were marginally more likely to be infected with *Bartonella* in the last two years of our study compared to the first year (Fig. 2A). This finding only partially supported our first hypothesis that infections would be more prevalent in the smaller habitat fragment and in later years in response to increased fragmentation, as we did not observe differences in *Bartonella* infection prevalence between our larger and smaller fragments. The significant differences in infection between the first year of the study and the last years may reflect a potential lag in the effects of forest loss that may have both preceded our work and occurred in earlier years of our study. However, this possibility is difficult to explore since land use/land cover data were not available for the first two years of our study. Additionally, the difference in infection likelihood across sites in 2021 compared to other years may indicate an effect of seasonality, as 2021 sampling occurred in November instead of our typical sampling period during April and May. Despite site not predicting *Bartonella* infection likelihood and the rarity of site-switching bats in this dataset (n = 1) and as previously reported from a broader mark-recapture dataset of this population (Becker et al., 2021), we observed strong synchrony in *Bartonella* prevalence between sites across multiple annual lags and in different directions (Fig. 2C), which may be driven by encounters between vampire bats from different sites during foraging (Greenhall et al., 1971). Synchrony between sites suggests that forest clearing may not be enough to truly isolate vampire bats in the region due to the affinity for feeding on livestock, which are common in the matrix but not within the LAR or KK. Indeed, previous work has shown that over time, the diets of vampire bats from the LAR and KK have converged, with stable isotope data indicating a shift from a more heterogeneous diet to one primarily consisting of livestock blood (Becker et al., 2018b).

While site did not predict *Bartonella* infection likelihood, the interaction between site and year was an important predictor of hemoplasma infection. Indeed, the sharp peak in hemoplasma prevalence that we observed in KK in 2017 coincided with marginally significant differences between infection likelihood in KK both before and after 2017, and in the LAR in the years following this peak, starting in 2019. The differences in infection likelihood in KK across years, in addition to the lack of significant differences in infection likelihood in the LAR over time, suggest that infection prevalence is more stable in the LAR than in KK (Fig. 2B). Further, while hemoplasma infection trends in KK appear to be influenced by those in the LAR, those in the LAR seem to be unaffected by infections in KK. We found a high degree of positive synchrony in infection likelihood between KK and the LAR at the negative two-year lag (Fig. 2D), indicating that hemoplasma infection trends in the LAR tend to lead those in KK by two years.

Much more diversity was observed in *Bartonella* compared to hemoplasmas, although this may be due to different phylogenetic similarity thresholds for delineating *Bartonella* (i.e. 96% similarity of the *gltA* gene; La Scola et al., 2003; Becker et al., 2018a) and hemoplasma genotypes (i.e. 98.5% similarity of the 16S rRNA gene; Becker et al. 2020b). While we observed a total of 16 *Bartonella* genotypes throughout this study, five main genotypes dominated (n > 20 each while n < 10 for all others; Fig S2). Three of these main genotypes (DR8, DR9, and DR11) showed significant differences in frequency among years, suggesting that some genotypes (i.e. DR1 and DR2) may be better established and less influenced by extrinsic factors than others (Fig. S3). Spatiotemporal factors were not predictors of infection likelihood for the main genotypes despite their frequencies differing significantly during a two year period from 2018– 2019, and in 2016 when only DR8 was detected.

While hemoplasma genotype richness was much lower than that seen in *Bartonella*, we observed complex dynamics at the genotype level in hemoplasmas. In keeping with previous studies of this system (Volokhov et al., 2017; Becker et al., 2020b, DeAnglis et al., 2024), we identified a total of five genotypes, two of which were only observed once. These latter genotypes were previously observed only in non-vampire bat species (i.e. *Neoeptesicus furinalis, Saccopteryx bilineata, Glossophaga soricina, Molossus nigricans,* and *Molossus alvarezi*), suggesting limited but existing possibility for transmission between sympatric bat species.

Additionally, the 17 vampire bats found to be coinfected with multiple genotypes may indicate a robust contact network between individuals that facilitates the transmission of hemoplasmas within the LAR, KK, and the surrounding area.

All three of the main hemoplasma genotypes differed in frequency over time but apparently reached equilibrium in the two most recent years of the study. While the frequency of VB1, the most commonly observed genotype, decreased from 2017–2019, that of VB2, the second most common genotype, increased during the same timespan. Meanwhile, VB3 frequency steadily increased throughout the entire study period (Fig. 7). Together, these trends suggest that competition among genotypes, especially between VB1 and VB2, may have resulted in a more balanced distribution in the hemoplasma community within vampire bats in recent years and allowed for the rise of VB3 (Mideo et al., 2008). While the infection status of hemoplasmas as a whole were influenced by the interaction between site and year, that of VB1 and VB2 were best predicted by year alone, with differences in infection likelihood further supporting the respective changes in frequency we observed during 2017–2019. In contrast with the most abundant hemoplasma genotypes and hemoplasmas overall, we did not identify significant differences in VB3 likelihood between years despite year being a marginal predictor. Differences in infection dynamics between hemoplasmas as a whole, and specific genotypes, as well as between genotypes, suggests that different hemoplasmas vary in their within-host dynamics and transmission strategies (Mideo et al., 2008; Rohani & Handel, 2015).

### Responses to forest loss were variable between and within pathogens

While site and tree cover both influenced infection likelihoods, both strength and interaction effects differed between *Bartonella* and hemoplasmas. *Bartonella* prevalence was positively affected by decreasing tree cover in the matrix, but only within the LAR (Fig. 4), suggesting that tree cover loss in the region as a whole may lead to more *Bartonella* infections in the larger habitat patch but not the smaller habitat patch. It is possible that the decline in tree cover may be enhancing *Bartonella* transmission by facilitating contacts between bat flies and bats (Mello et al., 2023), by altering immunity and stress responses (Messina et al., 2018), or by forcing bats that lived in the destroyed forest to move in the LAR. While we expected that habitat fragmentation would increase infection likelihood in KK more so than in the LAR, this was not supported by our data. It is possible that KK may be too small for additional loss in tree cover to affect overall *Bartonella* infection dynamics. Of the five main *Bartonella* genotypes we identified, only two were affected by forest loss, with differing importance of site and post-hoc significance (Fig. 5). This result again suggests that genotypes likely differ in transmission-relevant traits (e.g., infectiousness, infectious periods, etc.) that shape how their dynamics respond to habitat fragmentation (Mideo et al., 2008; Handel & Rohani, 2015; McCallum et al., 2017).

Hemoplasma infection likelihood was marginally influenced by both site and tree cover. In contrast with *Bartonella,* we did not find support for an interaction between these variables for hemoplasmas. As expected, hemoplasma infection likelihood was marginally higher in KK than in the LAR, and, counter to our hypothesis, with increasing tree cover (Fig. 4). Because forest loss in this system has been found to be accompanied by an increase in rangeland (DeAnglis et al., 2024), it is likely that it also results in an increased presence of livestock. The expanded availability of the vampire bat’s preferred food source may allow for more investment in immunity that could allow bats to better clear or decrease susceptibility to hemoplasma infections (Becker et al., 2018b). Similar results have been observed in Eastern bluebirds, in which food supplementation was associated with increased antibody responses and decreased nestling parasitism (Knutie, 2020).

Two of the individual hemoplasma genotypes were more significantly affected by forest loss than hemoplasmas overall, but with opposing trends and differing importance of site (Fig. 6). The variation in responses to forest loss among pathogen genotypes may indicate differing modes of transmission between genotypes (Handel & Rohani, 2015), which may have been differentially affected by the loss of tree cover within the matrix. Additionally, as proposed to explain the spatiotemporal trends that we observed, there may be some degree of competition between genotypes, in which the decline in prevalence of some genotypes may allow for the increase of others (Mideo et al., 2008).

### Sex and reproductive status, but not age class, influenced infection dynamics

Despite their differences in responses to spatiotemporal factors and forest loss, *Bartonella* and hemoplasma infections were similarly influenced by sex and reproductive status. Our findings were in contrast with our second hypothesis, as we expected that reproduction would come at the cost of the ability to mount immune responses against bacterial infections, especially in females. We found that male bats were more likely to be infected with either pathogen than female bats (Fig. 3A-B). Possible drivers of this trend include the effects of testosterone on immunity and differences in behavior between sexes. It is typically assumed that testosterone should negatively affect immune function while positively affecting characteristics important for reproduction (Folstad & Karter, 1992; Muehlenbein & Bribiescas, 2005). However, other studies have found the relationship between testosterone and immunity to be more complex: increased testosterone levels may support innate immunity over adaptive immunity and lead to behaviors that enhance transmission (Ezenwa et al., 2012; Becker et al., 2018b). Indeed, the spread of these blood-borne infections may be facilitated by aggressive encounters that may draw blood. While bats of either sex engage in both friendly and aggressive encounters, males tend to be more aggressive and engage in fewer friendly interactions compared to females (Wilkinson, 1984; Wilkinson, 1985; Carter & Wilkinson, 2013; Carter & Leffer, 2015). Additionally, the *Bartonella* and hemoplasma genotypes identified here are not closely related to those found in livestock (Fig S2; Fig S7), suggesting that blood shared between bonded bats via regurgitation and feeding from prey are not drivers of infection.

In keeping with results from our previous work on hemoplasmas in vampire bats (Volokhov et al., 2017), reproductive bats were less likely to harbor hemoplasma infections than non-reproductive bats (Fig. 3D). This trend was also seen in our models of *Bartonella* infection likelihood, albeit with marginal significance (Fig. 3C). We found only one exception, where reproductive hemoplasma-infected bats were marginally more likely to be infected with VB2 than non-reproductive bats (Fig. 8C). These findings are surprising, as reproduction is classically expected to negatively impact immunity through life history trade-offs (Sheldon & Verhulst, 1996). However, studies in bats increasingly suggest that the relationship between reproduction and immunity may be more complex (Eskew et al., 2025). In some situations, trade-offs between immunity and reproductive functions may not occur (Ruoss et al., 2018), or reproduction may even support only certain aspects of immunity (Becker et al., 2018b). Counter to our expectations that infections should be more likely in subadult bats, neither *Bartonella* nor hemoplasmas were influenced by age class, suggesting that bats seem to maintain susceptibility throughout their life. However, we did observe exceptions to this trend; adult bats were marginally less likely to be infected with hemoplasma genotype VB1 (Fig. S8A) and *Bartonella* genotype DR1 but more likely to be infected with *Bartonella* genotype DR9 than subadults (Fig. S4A-B). These results, along with instances of infection and genotype switching among recaptured bats, suggest that adaptive immunity to *Bartonella and* hemoplasmas may be limited at best (Figs. S4 and S9). However, it is also possible that some bats switching from positive to negative infection status may be due to a reduction in bacterial loads to below levels detectable by PCR.

### IBT expectations vs actual results

Past applications of IBT to infectious disease dynamics in fragmented habitats suggest that pathogens should be more common in small, isolated habitats compared to habitats that are larger and more easily accessible (Reperant, 2010; Shaw et al., 2024). However, and in line with several previous studies (Faust et al., 2018; Becker et al., 2020a), the effects of habitat fragmentation on *Bartonella* and hemoplasma dynamics in vampire bats appear to be more complex. The low R^2^ values from most of our models suggest that additional factors beyond those that we evaluated may contribute to *Bartonella* and hemoplasma dynamics in these bats. Potentially important predictors may include edge effects of habitat fragmentation (Suzán et al., 2012), season (Field et al., 2015), ectoparasite loads and cycles (Espinal-Palomino et al., 2025; Stuckey et al., 2017), climatic drivers (Páez et al., 2017), and the presence of alternative hosts that may dilute or enhance pathogen transmission (Ostfeld & Keesing, 2000). Future studies with finer-scale temporal sampling of vampire bats, explicit assessments of alternative transmission routes, and consideration of both edge and host community effects could provide further insight into the transmission dynamics of these relevant bacterial pathogens under environmental change.

### Conclusions

While the influence of demographic variables on infection likelihood was broadly consistent between bartonellae and hemoplasmas, we found that the effect of habitat fragmentation on bacterial infection varied across both pathogens and genotypes. Our work highlights the role that within-pathogen variation can play in shaping population-level transmission dynamics (Mideo et al., 2008; Withenshaw et al., 2016; Becker et al., 2020b), especially in the context of environmental change. As similar results have been found in some other systems (Perrin et al., 2023), our findings add to a growing body of research that demonstrates the complexity of the ways in which habitat fragmentation can affect infectious disease dynamics in wildlife. Theories such as IBT are useful starting points for generating hypotheses about how anthropogenic change could impact the transmission of pathogens of interest, but such expectations should be explicitly tested before they can be applied to any management practices.

## Acknowledgements

For assistance with field logistics, bat sampling and permits, we thank Mark Howells, Neil Duncan, and staff of the Lamanai Field Research Center, as well as many colleagues who helped net bats during 2015–2022 research trips in Belize. We also thank Konstantin Chumakov for laboratory support during this study. This work was supported by the National Science Foundation (DEB 1601052), ARCS Foundation, Explorers Club, American Society of Mammalogists, American Museum of Natural History (Theodore Roosevelt Memorial Fund, Taxonomic Mammalogy Fund), National Geographic Society (NGS-55503R-19), and the Research Corporation for Science Advancement (RCSA). This work was conducted as part of Subaward No. 28365, part of a USDA Non-Assistance Cooperative Agreement with RCSA Federal Award No. 58–3022–0-005. DJB was also supported by the Edward Mallinckrodt, Jr. Foundation.

## Data Accessibility and Benefit-Sharing

Data are available in the Dryad Digital Repository: (https://doi.org/10.5061/dryad.02v6wwqg8).

## Author Contributions

D.J.B. and L.R.L. designed the study and interpreted results; L.R.L., K.E.D., and D.J.B. collected samples and performed lab work; D.V.V. performed lab work and data analysis; A.Y. assisted in data analysis; N.B.S. and M.B.F. coordinated fieldwork; L.R.L. analyzed the data and wrote the first draft of the manuscript; all authors reviewed and approved the manuscript.

## Notes

https://doi.org/10.5061/dryad.02v6wwqg8

### Competing Interest Statement

The authors have declared no competing interest.

