## Supplemental Tables and Figures for "Longitudinal impacts of habitat fragmentation on *Bartonella* and hemotropic *Mycoplasma* dynamics in vampire bats"

Table S1. Hemoplasma GenBank accession numbers for each gene sequenced.

| 16S | 23S | rpoB |
|---|---|---|
| KY932674.1 - KY932676.1, KY932680.1, KY932681.1, KY932685.1 - KY932700.1, KY932722.1, MH245119.1, MH245120.1, MH245123.1, MH245130.1, MH245167.1, MH245176.1- MH245181.1, MH245191.1 - MH245193.1, MK353807.1, MK353808.1, MK353815.1, MK353821.1, MK353828.1, MK353836.1, MK353839.1, MK353846.1, MK353863.1, MK353872.1, MK353880.1, MK353881.1, MK353886.1 - MK353888.1, OQ385153.1 - OQ385174.1, OQ533048.1, OQ546498.1 - OQ546569.1, OR783317.1 - OR783319.1 | OQ456393.1, OQ518926.1 - OQ518933.1, OQ518936.1 - OQ518942.1, OQ518945.1, OQ518946.1 | OQ554324.1, OQ554325.1, OQ554326.1, OQ554327.1 |

Table S2. Results of spatiotemporal GLMMs for *Bartonella* and hemoplasmas.

| | *Bartonella* | | Hemoplasmas | |
|---|---|---|---|---|
| Fixed effect | $\chi^2$ | *p* | $\chi^2$ | *p* |
| site | 1.75 | 0.19 | 0.01 | 0.91 |
| year | 23.2 | <0.01 | 3.76 | 0.71 |
| sex | 8.53 | <0.01 | 8.58 | <0.01 |
| reproductive status | 4.02 | 0.04 | 4.24 | 0.04 |
| age class | 0.35 | 0.55 | 1.75 | 0.19 |
| site*year | NA | NA | 13.8 | 0.02 |

Table S3. Results of tree cover GLMMs for *Bartonella* and hemoplasmas.

| | *Bartonella* | | Hemoplasmas | |
|---|---|---|---|---|
| Fixed effect | $\chi^2$ | *p* | $\chi^2$ | *p* |
| site | 2.01 | 0.16 | 2.71 | 0.10 |
| tree cover | 7.22 | <0.01 | 3.01 | 0.08 |
| sex | 10.49 | <0.01 | 7.35 | <0.01 |
| reproductive status | 3.04 | 0.08 | 6.74 | <0.01 |
| age class | 0.54 | 0.46 | 0.70 | 0.40 |
| site*tree cover | 4.12 | 0.04 | NA | NA |

Table S4. Results of spatiotemporal GLMMs for *Bartonella* genotypes.

| | DR1 | | DR2 | | DR8 | | DR9 | | DR11 | |
|---|---|---|---|---|---|---|---|---|---|---|
| Fixed effect | $\chi^2$ | *p* | $\chi^2$ | *p* | $\chi^2$ | *p* | $\chi^2$ | *p* | $\chi^2$ | *p* |
| site | 1.12 | 0.29 | 0.17 | 0.68 | 0.05 | 0.82 | 0.01 | 0.91 | 1.25 | 0.26 |
| year | 9.51 | 0.15 | 5.45 | 0.49 | 3.81 | 0.70 | 5.37 | 0.50 | 10.1 | 0.12 |
| sex | 0.30 | 0.59 | 0.75 | 0.39 | 2.97 | 0.08 | 1.26 | 0.26 | <0.01 | 0.93 |
| reproductive status | 0.07 | 0.80 | 0.03 | 0.87 | 1.44 | 0.23 | 0.17 | 0.68 | 0.61 | 0.43 |
| age class | 3.11 | 0.08 | 0.41 | 0.52 | 0.02 | 0.89 | 3.65 | 0.06 | 0.24 | 0.62 |

Table S5. Results of tree cover GLMMs for *Bartonella* genotypes.

| | DR1 | | DR2 | | DR8 | | DR9 | | DR11 | |
|---|---|---|---|---|---|---|---|---|---|---|
| Fixed effect | $\chi^2$ | *p* | $\chi^2$ | *p* | $\chi^2$ | *p* | $\chi^2$ | *p* | $\chi^2$ | *p* |
| site | 0.59 | 0.44 | <0.01 | 0.98 | 0.02 | 0.88 | 0.09 | 0.76 | 2.40 | 0.12 |
| tree cover | 6.69 | 0.01 | 1.51 | 0.22 | 1.44 | 0.23 | 0.13 | 0.72 | 0.94 | 0.33 |
| sex | 0.22 | 0.64 | 0.54 | 0.46 | 2.99 | 0.08 | 1.30 | 0.25 | 0.52 | 0.47 |
| reproductive status | 0.26 | 0.61 | 0.19 | 0.66 | 1.39 | 0.24 | 0.49 | 0.48 | 0.16 | 0.69 |
| age class | 2.07 | 0.15 | 0.05 | 0.82 | 0.02 | 0.90 | 3.54 | 0.06 | 0.01 | 0.92 |
| site*tree cover | NA | NA | NA | NA | NA | NA | 4.24 | 0.04 | NA | NA |

Table S6. Results of GLMs for number of *Bartonella* genotypes, infection status switching, and genotype switching for recaptured bats (n = 59).

| | Number of genotypes | | Infection status switching | | Genotype switching | |
|---|---|---|---|---|---|---|
| Fixed effect | $\chi^2$ | *p* | $\chi^2$ | *p* | $\chi^2$ | *p* |
| number of captures | 5.36 | 0.02 | 0.19 | 0.66 | 8.13 | <0.01 |
| sex | 1.06 | 0.30 | 1 | 0.32 | 1.93 | 0.17 |
| minimum age (years) | 1.02 | 0.31 | 0 | 1 | 0.81 | 0.37 |
| site | 0.27 | 0.60 | 0.13 | 0.71 | 0.69 | 0.40 |
| age class switching | 0.69 | 0.41 | 1.14 | 0.29 | 0.33 | 0.57 |

Table S7. Results of spatiotemporal GLMMs for hemoplasma genotypes.

| | VB1 | | VB2 | | VB3 | |
|---|---|---|---|---|---|---|
| Fixed effect | $\chi^2$ | $p$ | $\chi^2$ | $p$ | $\chi^2$ | $p$ |
| site | <0.01 | 0.93 | 0.78 | 0.38 | 0.56 | 0.45 |
| year | 23.1 | <0.01 | 18.8 | <0.01 | 11.5 | 0.07 |
| sex | 0.90 | 0.34 | 2.50 | 0.11 | <0.01 | 0.99 |
| reproductive status | 0.43 | 0.51 | 3.62 | 0.06 | 3.14 | 0.08 |
| age class | 3.86 | 0.05 | 1.25 | 0.26 | 1.50 | 0.22 |

Table S8. Results of tree cover GLMMs for hemoplasma genotypes.

| | VB1 | | VB2 | | VB3 | |
|---|---|---|---|---|---|---|
| Fixed effect | $\chi^2$ | $p$ | $\chi^2$ | $p$ | $\chi^2$ | $p$ |
| site | 0.23 | 0.63 | 0.01 | 0.91 | 0.08 | 0.78 |
| tree cover | 4.59 | 0.03 | 2.55 | 0.11 | 0.00 | 1.0 |
| sex | 0.06 | 0.81 | 0.62 | 0.43 | <0.01 | 0.95 |
| reproductive status | 0.03 | 0.87 | 1.50 | 0.22 | 2.46 | 0.12 |
| age class | 2.83 | 0.09 | 0.67 | 0.41 | 0.98 | 0.32 |
| site:tree cover | NA | NA | 7.17 | <0.01 | NA | NA |

Table S9. Results of GLMs for number of hemoplasma genotype, infection status switching, and genotype switching for recaptured bats (n = 59).

| | Number of genotypes | | Infection status switching | | Genotype switching | |
|---|---|---|---|---|---|---|
| Fixed effect | $\chi^2$ | $p$ | $\chi^2$ | $p$ | $\chi^2$ | $p$ |
| number of captures | 0.01 | 0.90 | 1.22 | 0.27 | 1.07 | 0.30 |
| sex | 0.03 | 0.86 | 0.09 | 0.77 | 0.08 | 0.78 |
| minimum age (years) | 0.04 | 0.85 | 0.28 | 0.60 | 0.32 | 0.57 |
| site | 0.01 | 0.91 | 0.31 | 0.58 | 1.97 | 0.16 |
| age class switching | 0.69 | 0.41 | 0.65 | 0.42 | <0.01 | 0.94 |

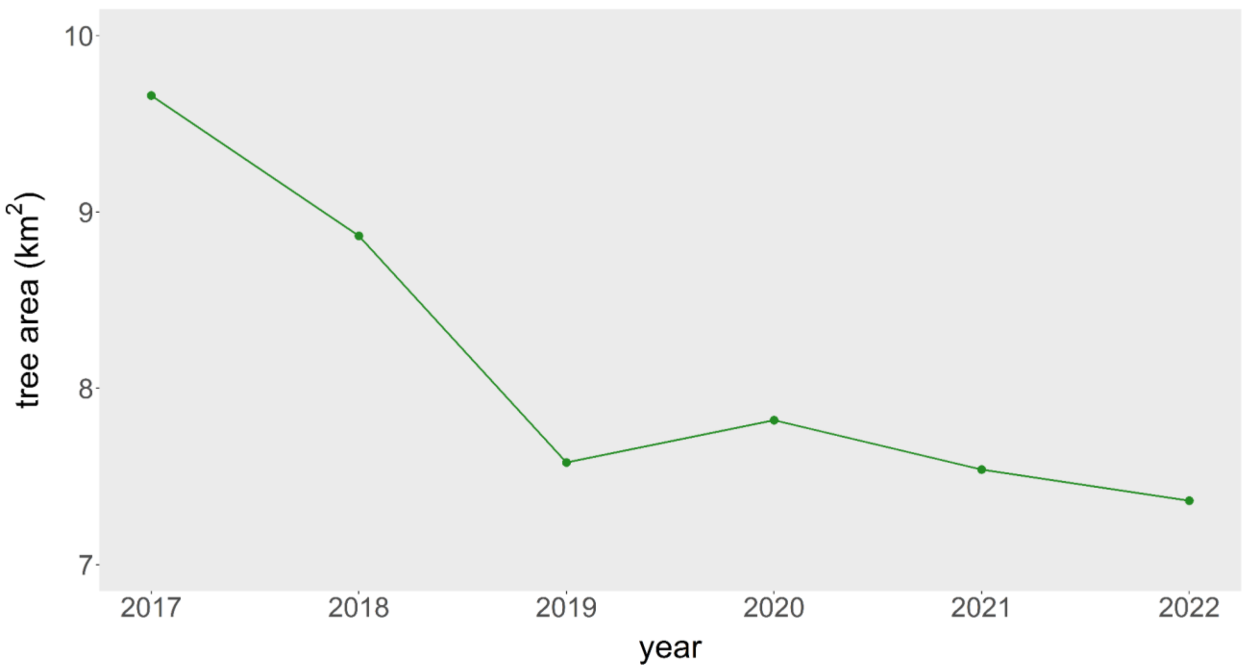


Figure S1. Change in tree cover in the matrix within 10 km of the central point between the LAR and KK from 2017-2022.

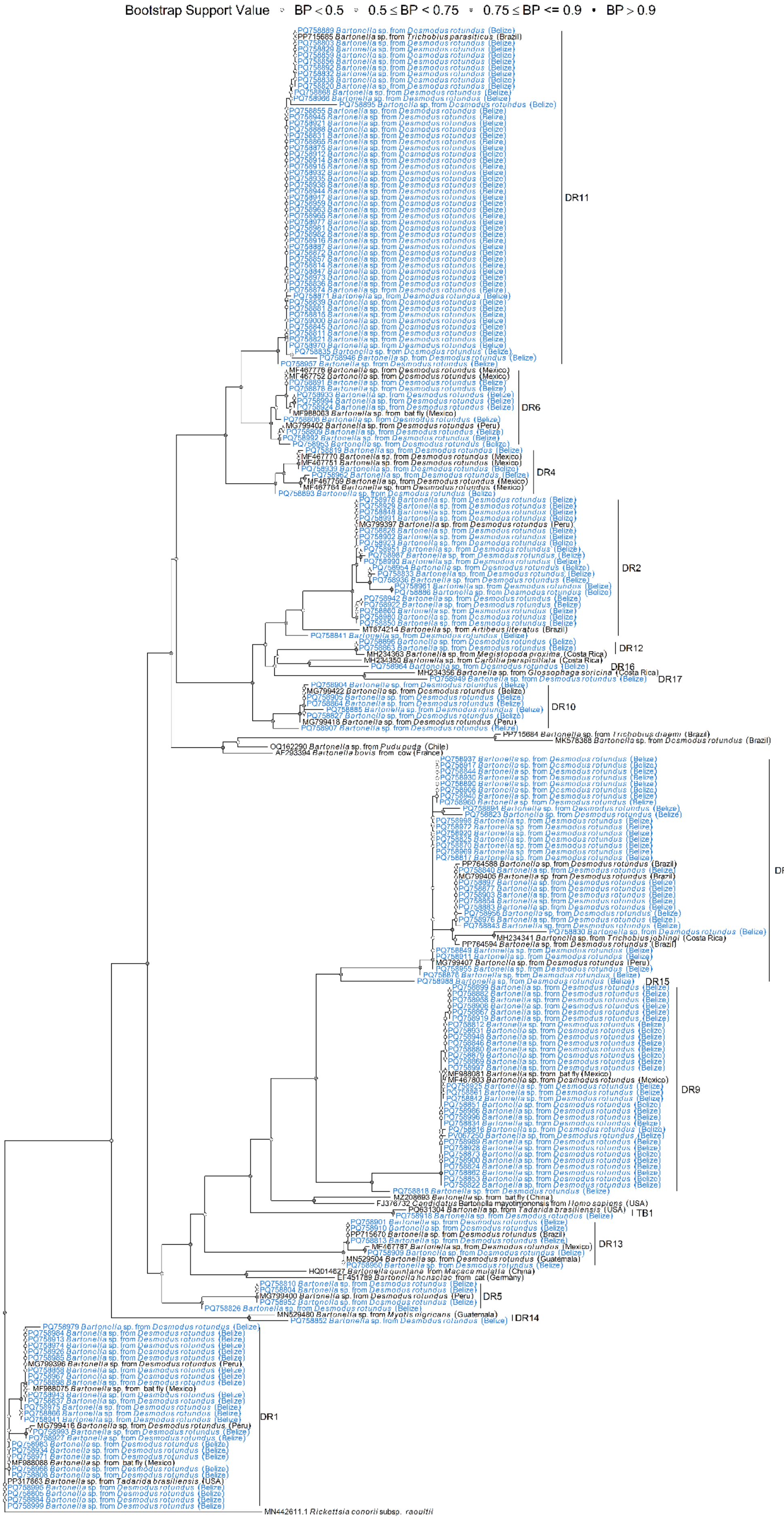


Supplementary Figure 2. Phylogenetic tree of *Bartonella* genotypes based on *gltA* gene sequence data constructed in NGPhylogeny.fr (Lemoine et al., 2019) using maximum likelihood with smart model selection (PhyML + SMS). A threshold of 96% similarly was used to guide genotype assignments. Sequences included in this analysis are shown in blue and top BLAST hits and other relevant *Bartonella* spp. are shown in black. Nodes are colored by bootstrap support value.

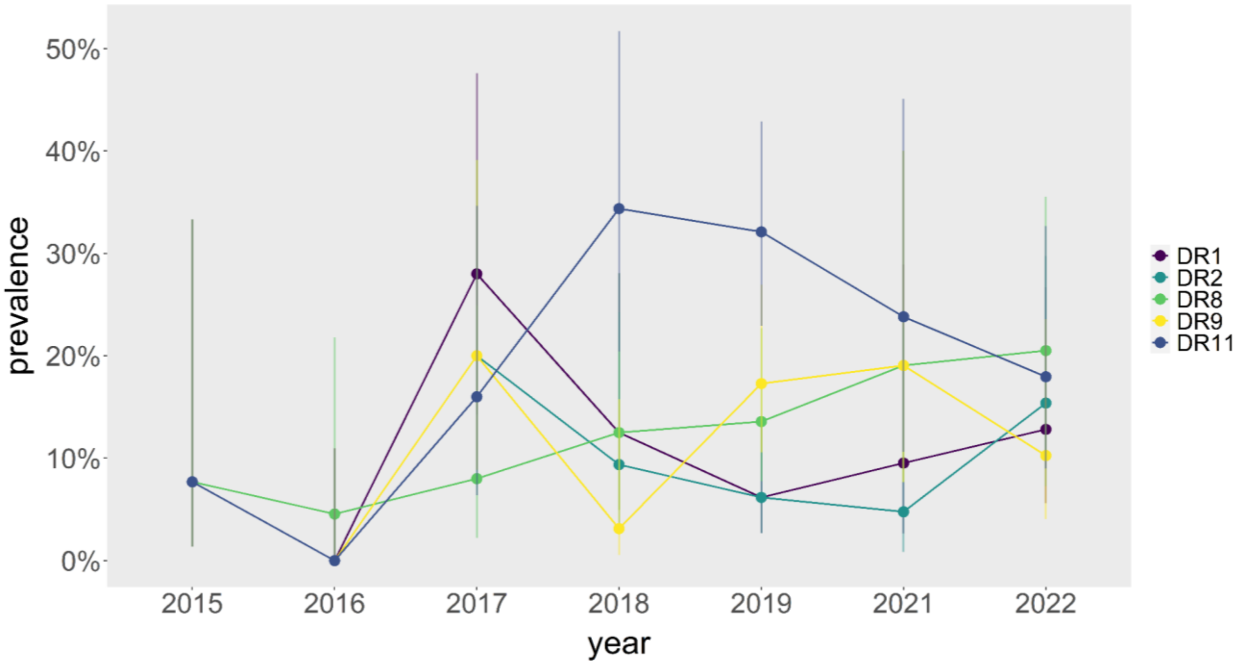


Figure S3. *Bartonella* genotype prevalence by year. Point estimates are shown with 95% confidence intervals (Wilson interval).

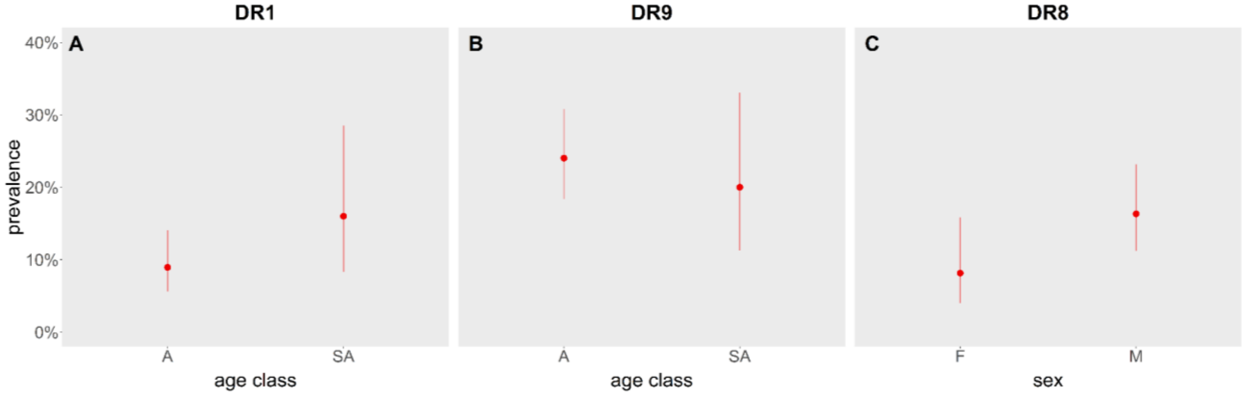


Figure S4. Infection prevalence of *Bartonella* genotypes DR1 (A) and DR9 (B) by age and DR8 by sex (C). Point estimates are shown with 95% confidence intervals (Wilson interval).

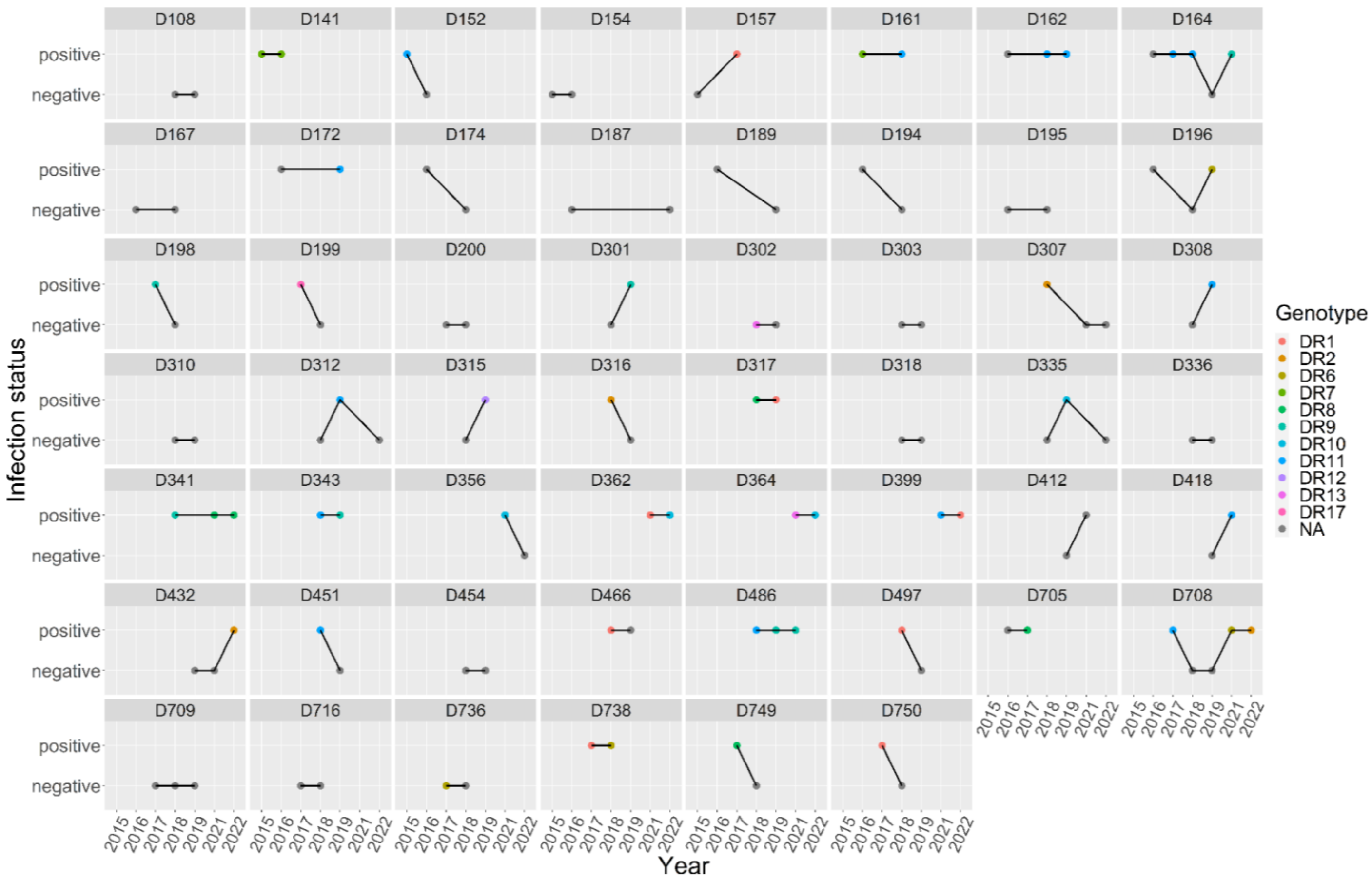


Figure S5. *Bartonella* infection status and genotypes over time in bats recaptured across multiple years.

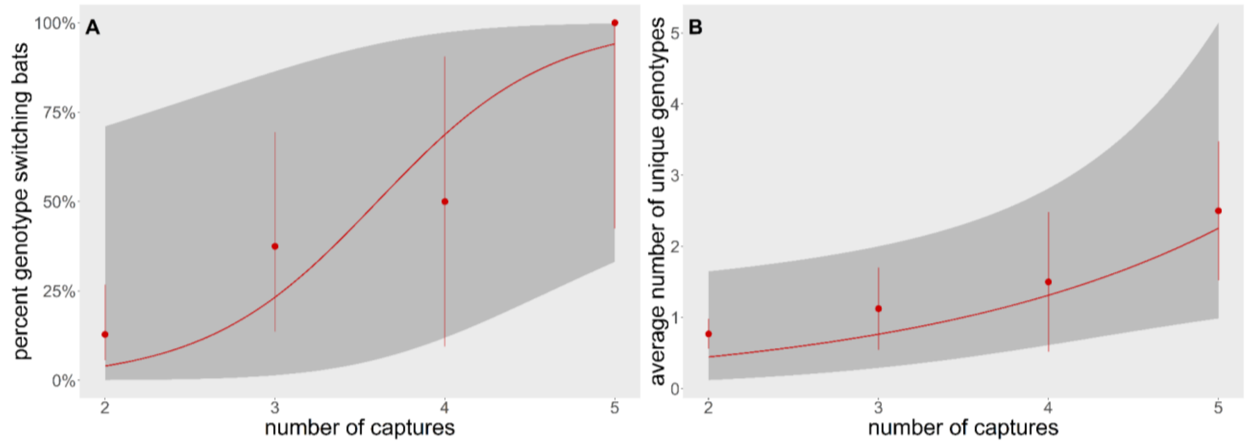


Figure S6. Total number of captures among recaptured vampire bats from 2015-2022 by the percent of bats that switched between genotype (A) and the average number of unique genotypes detected (B). Point estimates are shown with 95% confidence intervals.

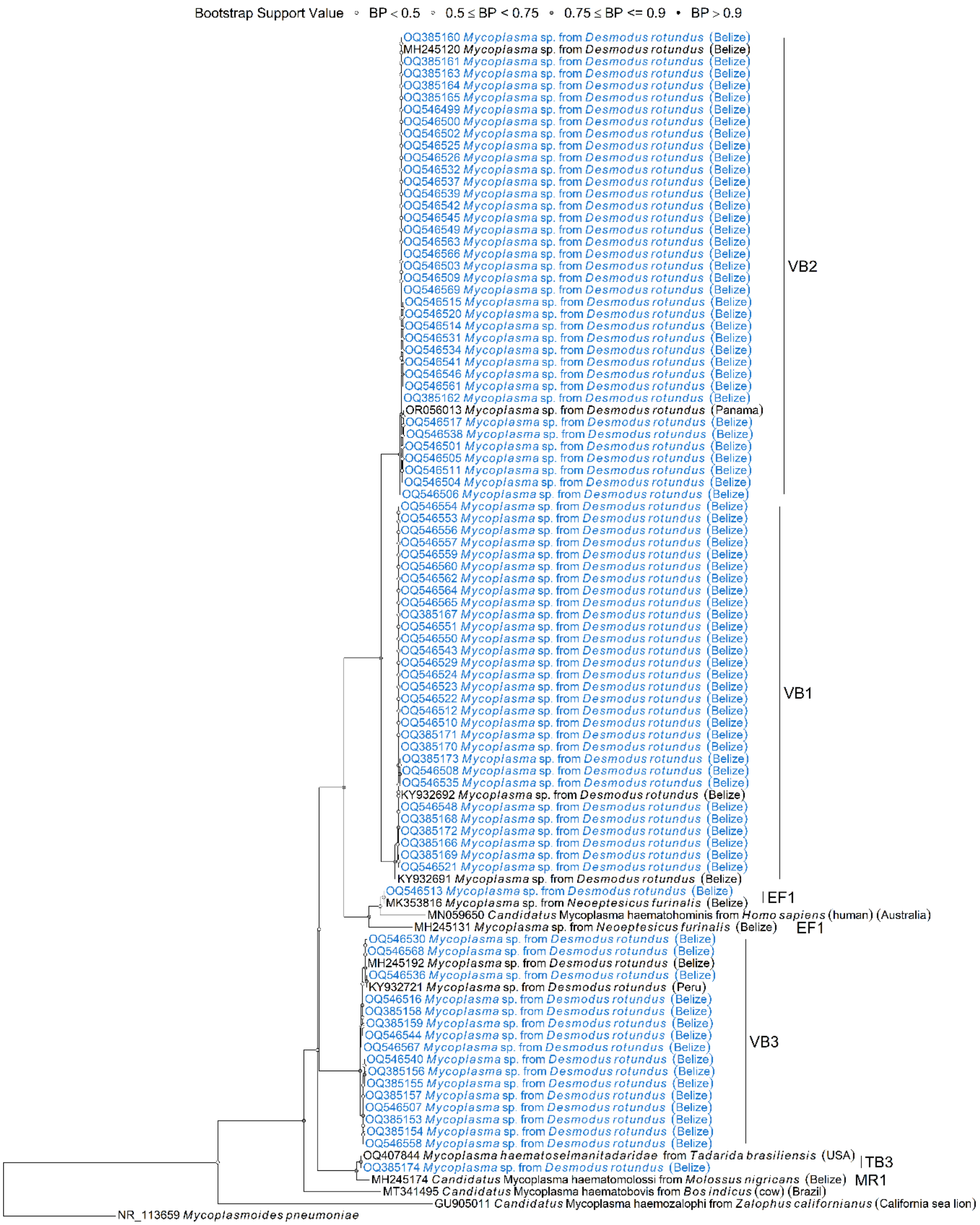


Figure S7. Phylogenetic tree of hemoplasma genotypes based on 16S rRNA gene sequence data constructed in NGPhylogeny.fr (Lemoine et al., 2019) using maximum likelihood with smart model selection (PhyML + SMS). A threshold of 98.5% similarly was used to guide genotypes assignments. Sequences included in this analysis are shown in blue and top BLAST hits and other relevant *Bartonella* spp. are shown in black. Nodes are colored by bootstrap support value.

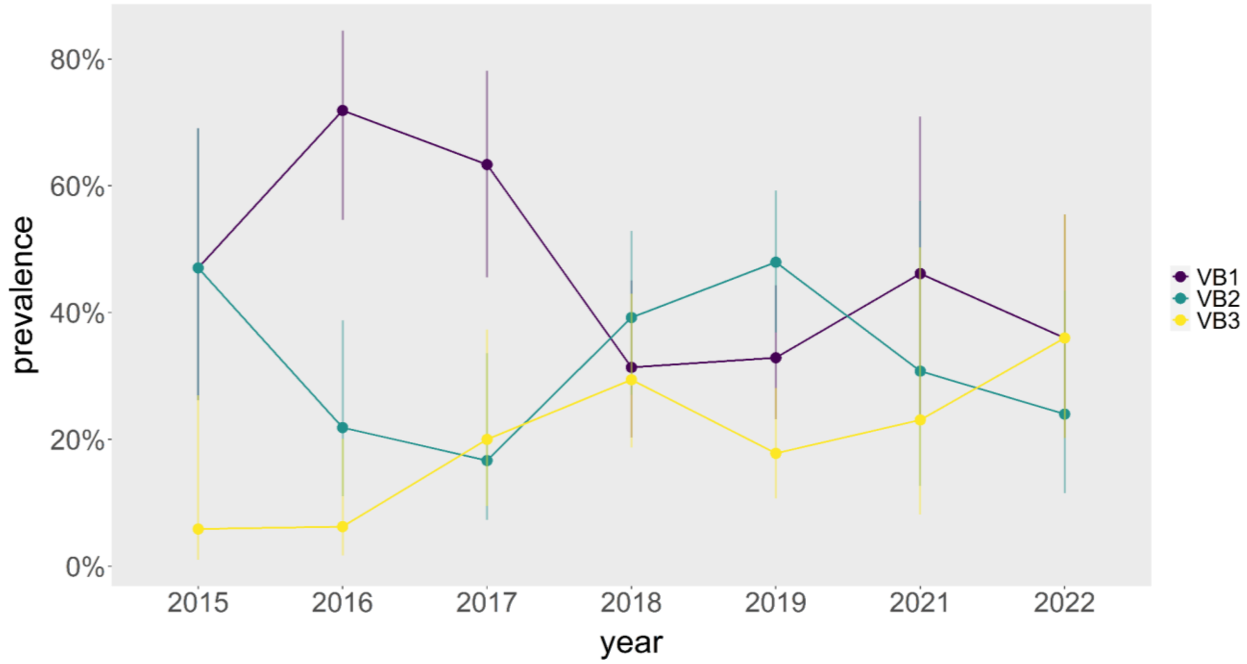


Figure S8. Prevalence of each hemoplasma genotype by year. Point estimates are shown with 95% confidence intervals (Wilson interval).

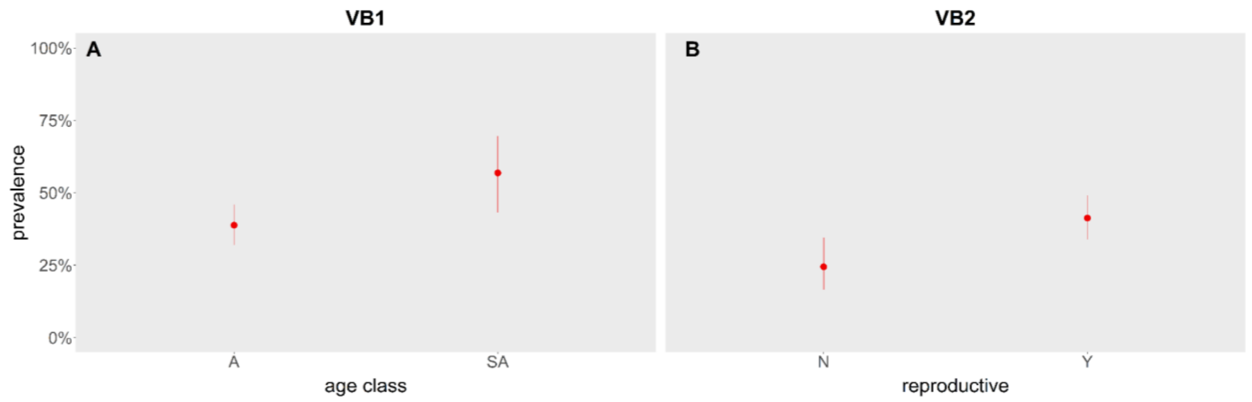


Figure S9. Prevalence of hemoplasma genotype A) VB1 by age class and B) VB2 by reproductive status. Point estimates are shown with 95% confidence intervals (Wilson interval).

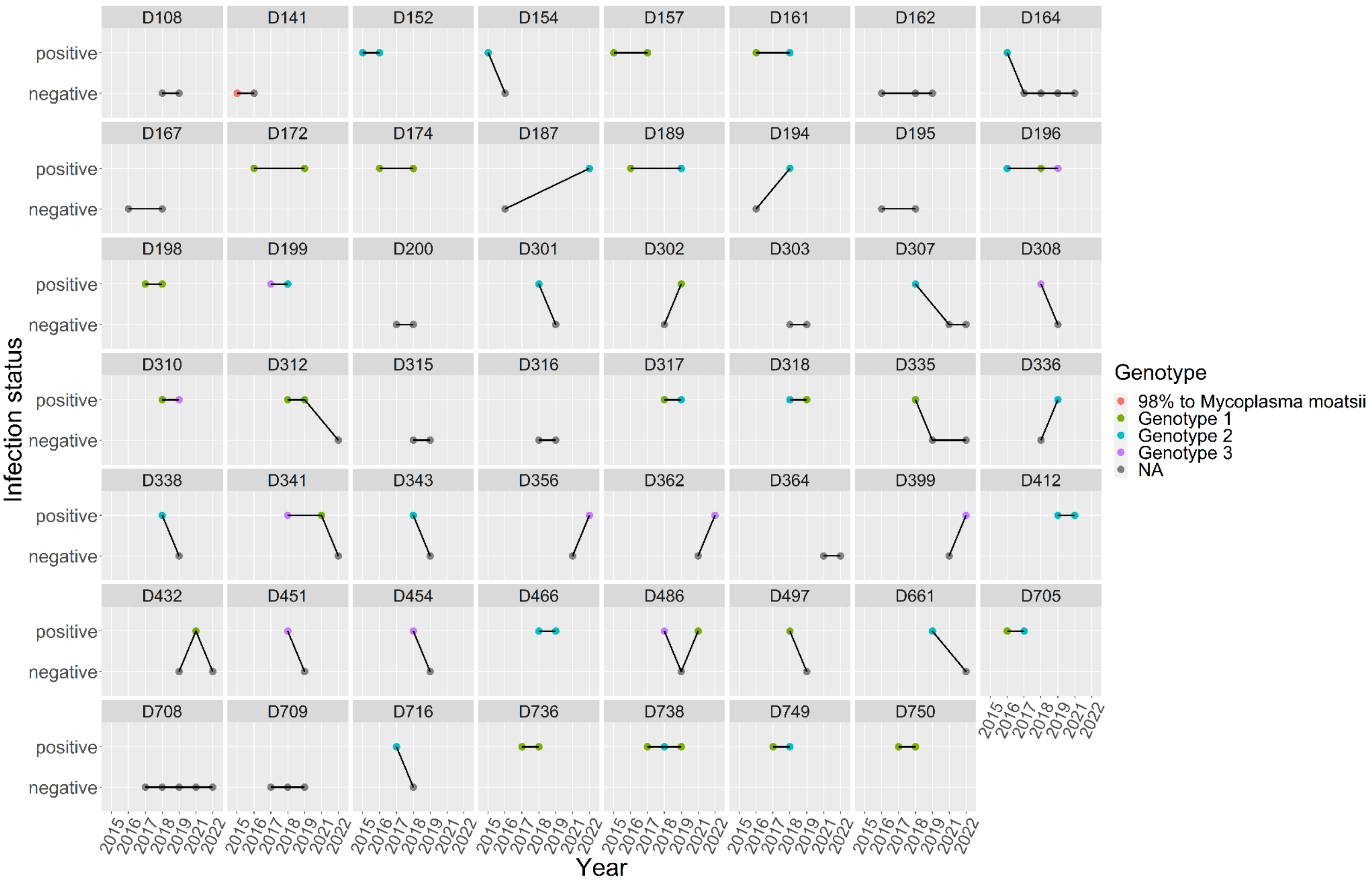


Figure S10. Hemoplasma infection status and genotypes over time in bats recaptured across multiple years.
